# Sequential infection reprograms the immune landscape to shape future responses

**DOI:** 10.64898/2026.06.26.734836

**Authors:** Mengdi Guo, Yanis Hichem Bouzaher, Diala Abd-Rabbo, Rene Quevedo, Heidi J. Elsaesser, Wenxi Xu, Melissa Yi Ran Liu, Fauzia N. Izzati, M Teresa Ciudad, Matthew Bianca, Zhe Qi Liu, Joao E.M de Oliveira, Arthur Mortha, Landon J. Edgar, Tracy L. McGaha, Tiffany A. Reese, David G Brooks

**Affiliations:** Princess Margaret Cancer Centre, University Health Network, Toronto, ON, Canada; Department of Immunology, University of Toronto, Toronto, ON, Canada; Department of Pharmacology and Toxicology, University of Toronto, Toronto, ON, Canada; Keenan Research Centre for Biomedical Sciences, St. Michael’s Hospital, Toronto, ON, Canada; Department of Immunology, University of Texas Southwestern Medical Center, Dallas, TX, USA; Department of Microbiology, University of Texas Southwestern Medical Center, Dallas, TX, USA

## Abstract

Mouse models have been instrumental in defining immune mechanisms but often fail to capture the complexity of human immunity, limiting clinical translation. A major limitation is the immunological immaturity of specific pathogen-free (SPF) mice relative to pathogen-experienced adult humans. Here, we use a sequential infection (SI) model that recapitulates cumulative pathogen exposure and define its impact on immune composition and function. Beyond the previously reported expansion of memory T cells, SI induced durable, system-wide remodeling across lymphoid and non-lymphoid tissues, reshaping innate and adaptive immune populations, tissue-resident immunity, and hematopoietic output. Single-cell transcriptomic analyses revealed inflammatory imprinting of naïve CD4 and CD8 T cells, whereas memory T cells acquired enhanced effector programs coupled with reduced biosynthetic activity, transcriptional states that more closely resemble those of pathogen-experienced adult humans. Functionally, SI mice recapitulated the human response to anti-CD28 super-agonist and exhibited altered magnitude and differentiation of acute and chronic antiviral T cell responses, demonstrating that cumulative pathogen exposure reshapes both existing immunity and the generation of future immune responses. Thus, cumulative pathogen exposure coordinately remodels hematopoiesis and naïve and memory lymphocyte states, establishing a durable inflammation-experienced immune landscape that reshapes both immune memory and future immune responses, with broad implications for the translational fidelity of preclinical mouse models.

## INTRODUCTION

Laboratory mice housed under specific pathogen-free (SPF) conditions are fundamental for elucidating immune mechanisms and developing life-saving therapies. Yet, despite sharing substantial genetic homology with humans (1), they often fail to recapitulate human immune responses (2, 3), contributing to numerous therapies that succeed preclinically but fail in the clinic with major economic and safety consequences (4, 5). Many factors likely contribute to these translational failures, including diet, housing conditions, and age. However, a major limitation is the immunological immaturity of the SPF immune system.

From birth, the human immune system is continually shaped by infections, vaccinations, and environmental exposures that drive progressive maturation and durable immune adaptation. In contrast, SPF mice are raised under conditions that markedly restrict antigenic experience, limiting the development of immunological memory, trained immunity, and hematopoietic remodeling. Although SPF mice encounter a restricted set of microbes, these exposures are insufficient to reproduce the diversity of stimuli that educate the human immune system. Consequently, SPF mice remain immunologically immature and more closely resemble neonatal humans than pathogen-experienced adults (6–10).

In 2016, Masopust and colleagues demonstrated that compared to SPF mice, feral and pet-store mice possess immune systems that closely resemble adult humans, including enhanced lymphocyte memory differentiation, tissue homing, and distinct transcriptional programs (7). Co-housing SPF mice with pet-store mice transferred this adult-like immune phenotype and altered responses to bacterial, parasitic, and viral infections (7). These results established that environmental immune experience has durable functional consequences and inspired the development of additional “dirty mouse” models, including rewilding in outdoor enclosures (11), and fetal transfer of SPF embryos into wild mice (8). However, widespread implementation of these approaches is limited by specialized infrastructure, increased genetic variability, and uncontrolled environmental exposures (6, 10). To overcome these limitations, Reese and colleagues developed a sequential infection (SI) model in which SPF mice are exposed to defined biosafety level 2 pathogens beginning at weaning (12). SI mice acquire many hallmarks of immune maturation observed in pet-store mice and adult humans, including expanded memory differentiation, adult-like interferon-associated transcriptional programs, and altered yellow fever specific vaccine responses, while maintaining experimental control (12).

Despite the growing use of sequential infection models, the mechanisms by which cumulative pathogen exposure remodels immune homeostasis and shapes future immune responses remain poorly understood. Here, we show that sequential infection reshapes the composition and function of adaptive and innate immunity, reprograms naïve lymphocytes toward an inflammation-imprinted state, and remodels hematopoiesis, collectively altering the differentiation of future immune responses. These changes produce adult human-like responses to immunological perturbation, including heightened sensitivity to CD28 super-agonist and altered CD8 T cell differentiation during acute and chronic viral infection. Thus, immune history is a fundamental determinant of immune homeostasis, revealing how cumulative pathogen exposure establishes a durable immune landscape that shapes both existing memory and the generation of subsequent immune responses.

## RESULTS

### Sequential infection drives immune remodeling in the peripheral blood and secondary lymphoid organs

We established a sequential infection (SI) protocol based on Reese *et al.* (12), exposing SPF mice beginning at 4 weeks of age to murine gammaherpesvirus 68 (MHV68; a model of Epstein–Barr virus and Kaposi’s sarcoma-associated herpesvirus, intranasal), murine cytomegalovirus (MCMV; a model of human CMV, intraperitoneal), influenza A (H1N1 A/Puerto Rico/8/1934, intranasal), and *Citrobacter rodentium* (a model of enteropathogenic *E. coli*, oral gavage) at two-week intervals (Fig. 1A, Table S1). Mice were rested for 8 weeks after the final infection, and age-matched mock-infected SPF (miSPF) mice served as controls. Experiments were performed at 18 weeks of age, corresponding approximately to a 25-year-old human adult (13). By combining pathogens with distinct tissue tropisms, infection routes, and persistence profiles, this model was designed to recapitulate cumulative immune experience during early life.

**Figure 1.**
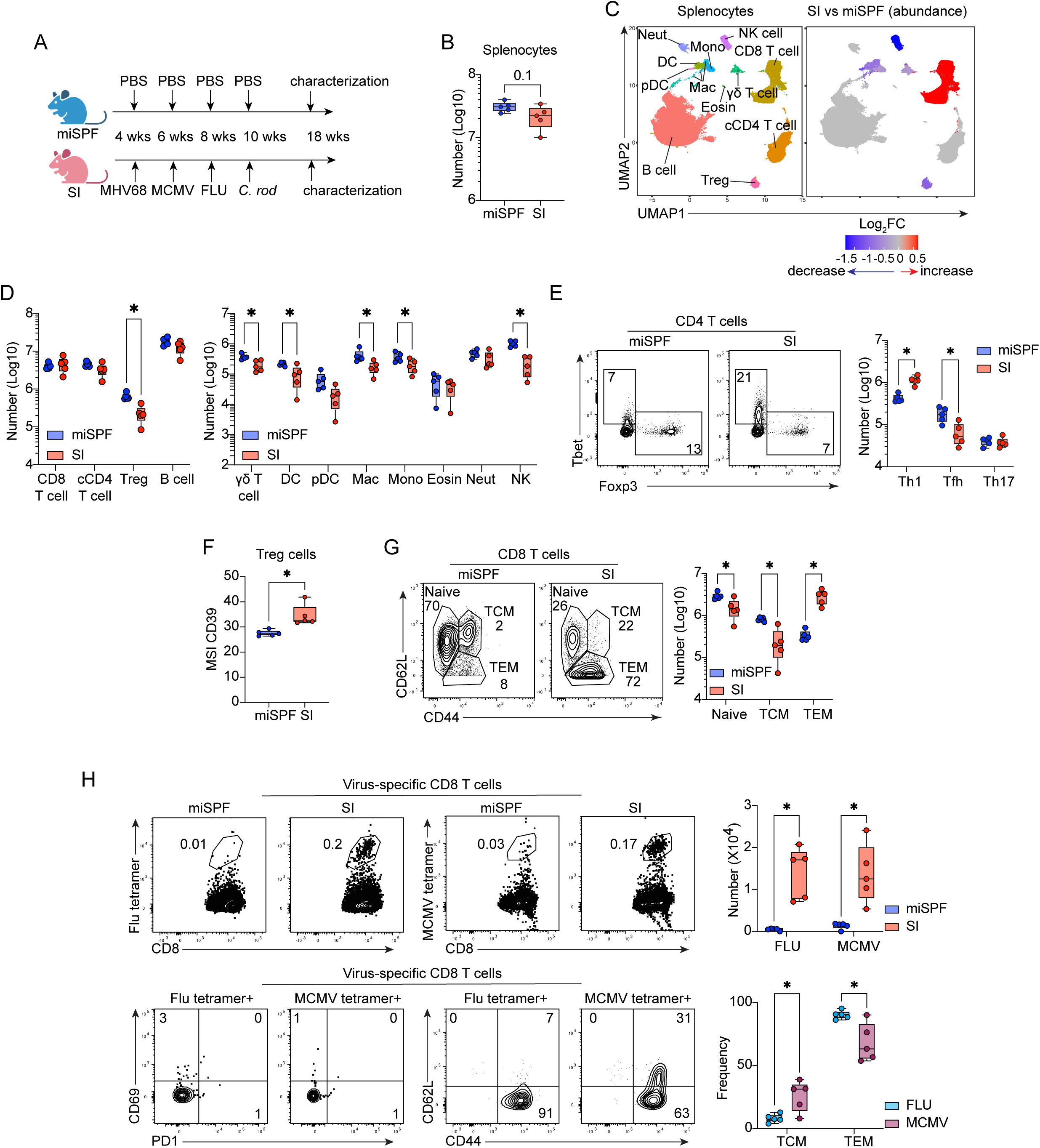
Sequential infection remodels immune composition and promotes T cell memory differentiation in the spleen. **(A)** Schematic of the SI regimen. **(B–H)** Immune composition of spleens from 18-week-old miSPF and SI mice characterized by CyTOF (**C, D, G**) and flow cytometry (**E, F, H**). **(B)** Total number of splenocytes. **(C)** UMAP visualization of splenic immune subsets identified by Phenograph clustering. (Left) All immune populations. (Right) Populations with significant changes in abundance, determined using diffcyt. Red: increased in SI mice; blue: decreased in SI mice; gray: unchanged. **(D)** Number of the indicated immune subsets. **(E, F)** Number and phenotype of CD4 T helper (Th) subsets: Tbet^+^ Th1, Bcl6^+^ Tfh, and RORγt^+^ Th17 cells, and FoxP3^+^ Treg. (G) Representative flow plots showing CD8 naïve, TCM, and TEM CD8 cells, with quantification of each subset. (H) Frequency and number of influenza (NP_366–374_)- and MCMV (M45_985-993_)-specific CD8 T cells (top), and phenotypic distribution of virus-specific CD8 T cell subsets (bottom). Neut, neutrophil; Mac, macrophage; Alv Mac, alveolar macrophage; Mono, monocyte; Eosin, eosinophil; pDC, plasmacytoid dendritic cell. Data are pooled from three independent experiments (≥4 mice per group). Circles represent individual mice. In box-and-whisker plots, boxes indicate the interquartile range, center lines indicate the mean, and whiskers denote the minimum and maximum values. Statistical significance was determined using an unpaired two-tailed t test. *, p<0.05.

Sequential infection induced broad remodeling of the peripheral blood compartment. Compared with miSPF controls, SI mice exhibited reduced circulating NK cells, granulocytes, monocytes, and dendritic cells, together with a relative expansion of T cells and increased plasma IgG levels (Fig. S1A, S1B), consistent with environmentally exposed “dirty” and rewilded mice (7, 14, 15). Unlike co-housed and rewilded mice, SI mice did not develop granulocytosis, likely reflecting the absence of chronic fungal colonization in this controlled model (14, 16, 17). Thus, SI recapitulates key hallmarks of environmentally acquired immune maturation while maintaining experimental control.

Within the spleen, total cellularity was largely preserved despite a trend toward reduced cell numbers (Fig. 1B), indicating that SI primarily alters immune composition rather than tissue size. SI mice exhibited reduced numbers of Tregs, γδ T cells, dendritic cells, macrophages, monocytes, and NK cells (Fig. 1C, D, S1C). Macrophages upregulated MHCII and downregulated CD86, whereas dendritic cells maintained stable expression of MHCII, CD80, and CD86 (Fig. S1D), consistent with observations in pet-store mice (18). SI mice also displayed reduced CD62L⁺ NK cells (Fig. S1E), suggesting that prior pathogen exposure promotes differentiation of NK cells with enhanced tissue-homing potential (19). Thus, the sequential infections induce durable remodeling of innate immune composition and activation state.

Although total cCD4 T cell numbers were unchanged (Fig. 1D), SI markedly altered their differentiation. Effector memory (TEM, PD1⁻CD44⁺CD62L⁻) CD4 T cells expanded at the expense of central memory (TCM, PD1⁻CD44⁺CD62L⁺) and naïve (PD1⁻CD44⁻CD62L⁺) populations in SI mice (Fig. S1F), consistent with progressive antigen-driven maturation (20). This shift was accompanied by increased T-bet⁺ Th1 cells and reduced Bcl6⁺ Tfh and FoxP3⁺ Treg populations (Fig. 1D, E), indicating polarization toward inflammatory effector programs. Interestingly, the remaining Tregs expressed higher levels of CD39 (Fig. 1F), suggesting enhanced suppressive capacity (21). Thus, sequential infection not only expands immune memory but also qualitatively reshapes CD4 T cell function.

A similar pattern was observed in the CD8 compartment. Total CD8 T cell numbers remained stable despite a modest increase in relative abundance (Fig. 1C, D), whereas differentiation shifted toward TEM cells with reciprocal reductions in naïve and TCM populations (Fig. 1G). These changes closely resemble the CD8 T cell architecture of dirty mice and adult humans (7, 16, 18, 22, 23), indicating sustained maturation of the cytotoxic T cell compartment. Influenza (NP_366–374_)-specific and MCMV (M45_985-993_)-specific CD8 T cells were readily detected in SI but not miSPF mice (Fig. 1H). These cells virus-specific CD8 T cells consisted predominantly of TEM cells with smaller TCM populations and lacked markers of recent activation (CD69, PD-1), consistent with stable long-term memory (Fig 1H). MCMV-specific cells were relatively enriched for TCM compared with influenza-specific cells, paralleling memory differentiation patterns observed in human CMV and influenza infection (24). Similar shifts toward memory differentiation were observed in mediastinal, inguinal, and mesenteric lymph nodes despite unchanged cellularity (Fig. S1G, S1H), demonstrating that sequential infection establishes a stable antigen-experienced T cell compartment throughout secondary lymphoid organs.

### Sequential infection drives tissue-specific immune remodeling and resident memory formation in non-lymphoid organs

We next examined how sequential infection reshapes immune composition within non-lymphoid tissues. Using *in vivo* antibody labeling to distinguish circulating from tissue-infiltrating leukocytes (25), we observed a ∼6-fold increase in lung-infiltrating immune cells in SI mice compared with miSPF controls (Fig. 2A). This expansion was accompanied by broad remodeling of the tissue immune compartment (Fig. 2B, S2A) and was driven primarily by adaptive immune populations, including CD8 and CD4 T cells, Tregs, γδ T cells, and B cells, whereas neutrophils were the only innate population that increased substantially (Fig. 2C). Thus, cumulative pathogen exposure promotes the long-term establishment of immune populations within peripheral tissues rather than transient inflammatory infiltration.

**Figure 2.**
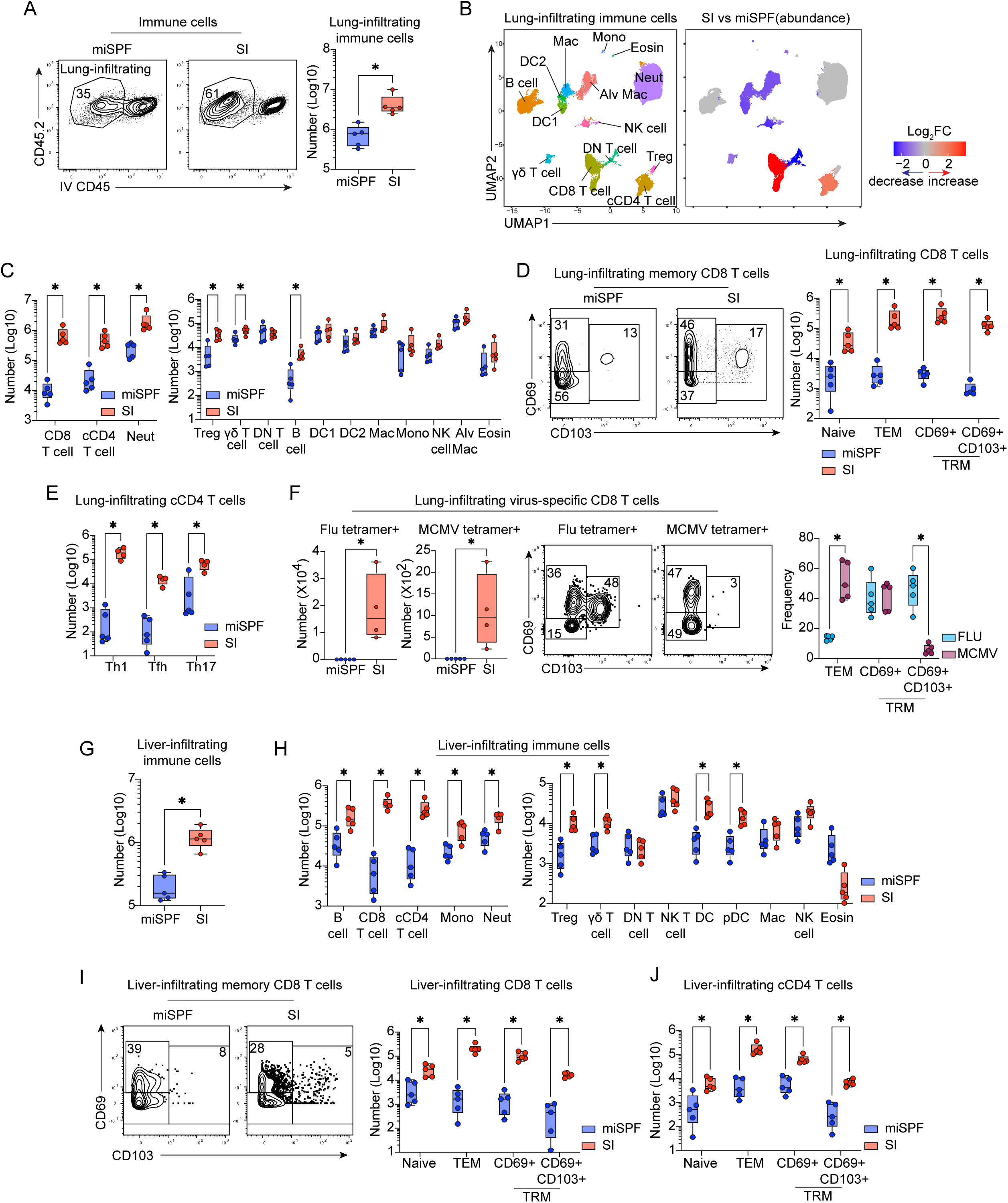
Sequential infection establishes mature tissue immune landscapes in the lung and liver. **(A–F)** Lung and **(G–J)** liver immune composition in 18-week-old miSPF and SI mice analyzed by CyTOF or flow cytometry. **(A)** Intravascular (i.v.) anti-CD45 labeling strategy used to distinguish circulating (CD45.2⁺ CD45⁻) from tissue-infiltrating (CD45.2⁺ CD45⁺) immune cells. Quantification of lung-infiltrating immune cells. **(B)** UMAP visualization of lung immune populations identified by Phenograph clustering. Left, all annotated populations. Right, populations exhibiting significant abundance changes determined by diffcyt. Red, increased in SI mice; blue, decreased in SI mice; gray, unchanged. **(C)** Number of lung-infiltrating immune subsets. **(D)** Representative flow plots and quantification of lung CD8 T cell subsets. **(E)** Number of lung-infiltrating CD4 Th subsets. **(F)** Number and phenotype of lung-infiltrating influenza-specific and MCMV-specific CD8 T cells. **(G)** Number of liver-infiltrating (IV-CD45⁻) immune cells. **(H)** Number of liver-infiltrating immune cells. **(I)** Representative flow plots and quantification of liver-infiltrating CD8 T cell subsets. **(J)** Number of liver-infiltrating CD4 T cell subsets. Data represent three independent experiments (≥4 mice/group). Circles represent individual mice. In box-and-whisker plots, boxes indicate the interquartile range, center lines indicate the mean, and whiskers denote the minimum and maximum values. Statistical significance was determined using an unpaired two-tailed t test. *, p<0.05.

The accumulation of adaptive immune cells was accompanied by marked changes in T cell differentiation. Lung CD8 T cells preferentially expanded as CD69⁺CD103⁺/⁻ tissue-resident memory (TRM) and TEM cells (Fig. 2D), closely resembling tissue-adapted memory populations observed in co-housed mice and adult humans (7, 23, 26). cCD4 T cells similarly expanded across TEM, TRM subsets and Th1, Tfh, Th17, and Treg compartments (Fig. 2C, 2E, S2B), indicating that sequential infection promotes both memory formation and functional diversification within peripheral tissues.

Although innate immune populations largely returned to baseline frequencies by 8 weeks after the final infection, evidence of persistent reprogramming remained. Lung NK cells were enriched for CD62L⁺ subsets (Fig. S2C), suggesting that the reduction of CD62L⁺ NK cells in lymphoid tissues reflects preferential recruitment into peripheral organs. Alveolar macrophages upregulated MHCII and downregulated CD80, consistent with durable innate adaptation distinct from that induced by single infections (27). Both macrophages and DCs increased PD-L1 expression, while DCs additionally upregulated CD40 and downregulated MHCII (Fig. S2D), collectively suggesting the establishment of a regulatory environment that may limit excessive tissue inflammation despite increased immune cell residency (28).

Antigen-specific CD8 T cells further highlighted the tissue-specific consequences of sequential infection. Lung influenza- and MCMV-specific CD8 T cells were readily detected in SI but not miSPF mice and exhibited a memory phenotype (CD44^hi^ PD1^dim^) (Fig. 2F, S2E). Influenza-specific cells were approximately 10-fold more abundant than MCMV-specific cells and were strongly biased toward CD69⁺CD103⁺ TRM differentiation, consistent with the lung tropism of influenza infection (Fig. 2F). In contrast, MCMV-specific CD8 T cells were distributed between TEM and CD69⁺ TRM populations with minimal CD103 expression (Fig. 2F), mirroring human memory responses (24). Thus, while cumulative pathogen exposure broadly promotes tissue residency, the organization of the memory compartment remains shaped by individual pathogen biology.

The liver exhibited a related but distinct pattern of immune remodeling. SI mice displayed increased immune infiltration (Fig. 2G) and expansion of T cells, B cells, Tregs, γδ T cells, DCs, pDCs, monocytes, and neutrophils (Fig. 2H, S2F, S2G). As in the lung, CD8 T cells were enriched for TEM and CD69⁺ TRM populations (Fig. 2I); however, CD103⁺ TRM cells were largely absent, consistent with liver-specific residency programs (29). Influenza- and MCMV-specific CD8 T cells similarly adopted CD69⁺CD103⁻ TRM and TEM phenotypes (Fig. S2H). cCD4 T cells expanded across memory and helper subsets (Fig. 2J, S2I), while Tregs increased in frequency and expressed elevated CD39 (Fig. 2H, S2J), suggesting redistribution and long-term maintenance within non-lymphoid tissues. Innate immune cells also exhibited evidence of durable adaptation, with DCs upregulating MHCII, CD80, CD39, and PD-L1 (Fig. S2K), and macrophages increasing MHCII expression (Fig. S2L).

In contrast to the lung and liver, total immune infiltration was not increased in the brain (Fig. S2M). However, the CD8 T cell compartment shifted from predominantly naïve to TEM- and TRM-enriched populations (Fig. S2N), indicating that sequential infection promotes immune maturation even within immune-privileged tissues. Thus, cumulative pathogen exposure drives durable tissue-specific immune remodeling, establishing mature resident immune populations and locally adapted immune niches throughout non-lymphoid organs.

### Sequential infection drives durable remodeling of the hematopoietic compartment

Infections can induce durable reprogramming of the bone marrow and alter long-term hematopoietic output (30–32). Given the widespread immune remodeling observed across peripheral tissues, we next asked whether cumulative pathogen exposure also reshapes hematopoiesis.

Compared with miSPF mice, SI mice exhibited reduced numbers of hematopoietic stem cells (HSCs), multipotent progenitors (MPPs), hematopoietic progenitors (HPC1 and HPC2), and common myeloid progenitors (CMPs) (Fig. S3A, S3B). The reduction in myeloid-biased progenitors, particularly CMPs, is consistent with the decreased abundance of monocytes and macrophages observed in the blood and secondary lymphoid organs (Fig. 1D, S1A), suggesting that peripheral immune remodeling is accompanied by altered cellular output from the bone marrow. In contrast, common lymphoid progenitors (CLPs) were largely preserved, mirroring the stable numbers of cCD4 and CD8 T cells in the spleen and lymph nodes (Fig. 1D, S1H). Thus, repeated inflammatory exposure extends the effects of sequential infection beyond mature immune populations to reshape hematopoietic stem and progenitor compartments, reprogramming the cellular foundation from which future immune responses are generated.

### Sequential infection induces durable immune remodeling across sexes

To determine whether SI-induced immune remodeling persists over time, we analyzed the lungs of SI and miSPF mice at 22 weeks of age, four weeks after our primary analysis. SI mice maintained elevated immune cell infiltration comparable to that observed at 18 weeks (Fig. S3C), indicating that tissue remodeling was sustained well beyond the resolution of infection. The expansion of CD8 TEM and CD69⁺ TRM populations also remained largely stable, with only a modest reduction in the CD69⁺CD103⁺ TRM subset that remained greater than 100-fold above miSPF mice (Fig. S3D). Importantly, this contraction did not diminish antigen-specific immunity, as influenza- and MCMV-specific CD8 T cell numbers were preserved (Fig. S3E). Similarly, increased cCD4 T cell accumulation and Th subset distribution were maintained between 18 and 22 weeks with slightly reduced Tfh (Fig. S3F), demonstrating the long-term stability of both CD4 and CD8 T cell remodeling. Beyond T cells, SI mice retained comparable numbers of B cells, NKT cells, eosinophils, and NK cells, while exhibiting modest increases in DCs, macrophages, monocytes, and neutrophils (Fig. S3G). The increases in innate cells were very modest in age-matched miSPF mice over the same period (Fig. S3G), suggesting less of an age-dependent effect. Together, these findings indicate that sequential infection establishes a durable tissue immune landscape rather than a transient post-infectious state.

We next assessed whether these effects were consistent across sexes by comparing 18-week-old male and female miSPF and SI mice. Both sexes exhibited comparable increases in lung immune infiltration following sequential infection (Fig. S3H). Female SI mice displayed a modestly greater expansion of CD8 TEM and CD69⁺ TRM populations than males, whereas CD69⁺CD103⁺ TRM cells were present at similar frequencies (Fig. S3I). Likewise, cCD4 T cells expanded in both sexes, with slightly greater accumulation in females (Fig. S3J). Despite these quantitative differences, the overall pattern of immune remodeling was similar between males and females. Thus, sequential infection induces a robust and durable program of immune remodeling that is largely conserved across sex.

### Sequential infections uncouple total memory accumulation from antigen-specific memory maintenance

We next asked how cumulative pathogen exposure shapes memory T cell differentiation relative to individual infections by comparing SI mice to mice infected with each pathogen alone (Fig. 3A). While sequential infection markedly expanded the overall memory compartment, its effects on individual antigen-specific responses were unexpectedly distinct. Within the spleen, SI mice exhibited increased numbers of Th1 cells compared with most single infections, although MHV68 alone was sufficient to induce a comparable Th1 response (Fig. S4A). Similarly, all infections except *C.rod* promoted splenic CD8 TEM differentiation, with MHV68 driving the largest increase among individual pathogens (Fig. S4B). However, SI mice exhibited the greatest overall expansion of CD8 TEM cells, indicating that each successive infection contributes incrementally to the accumulation of the total memory pool.

**Figure 3.**
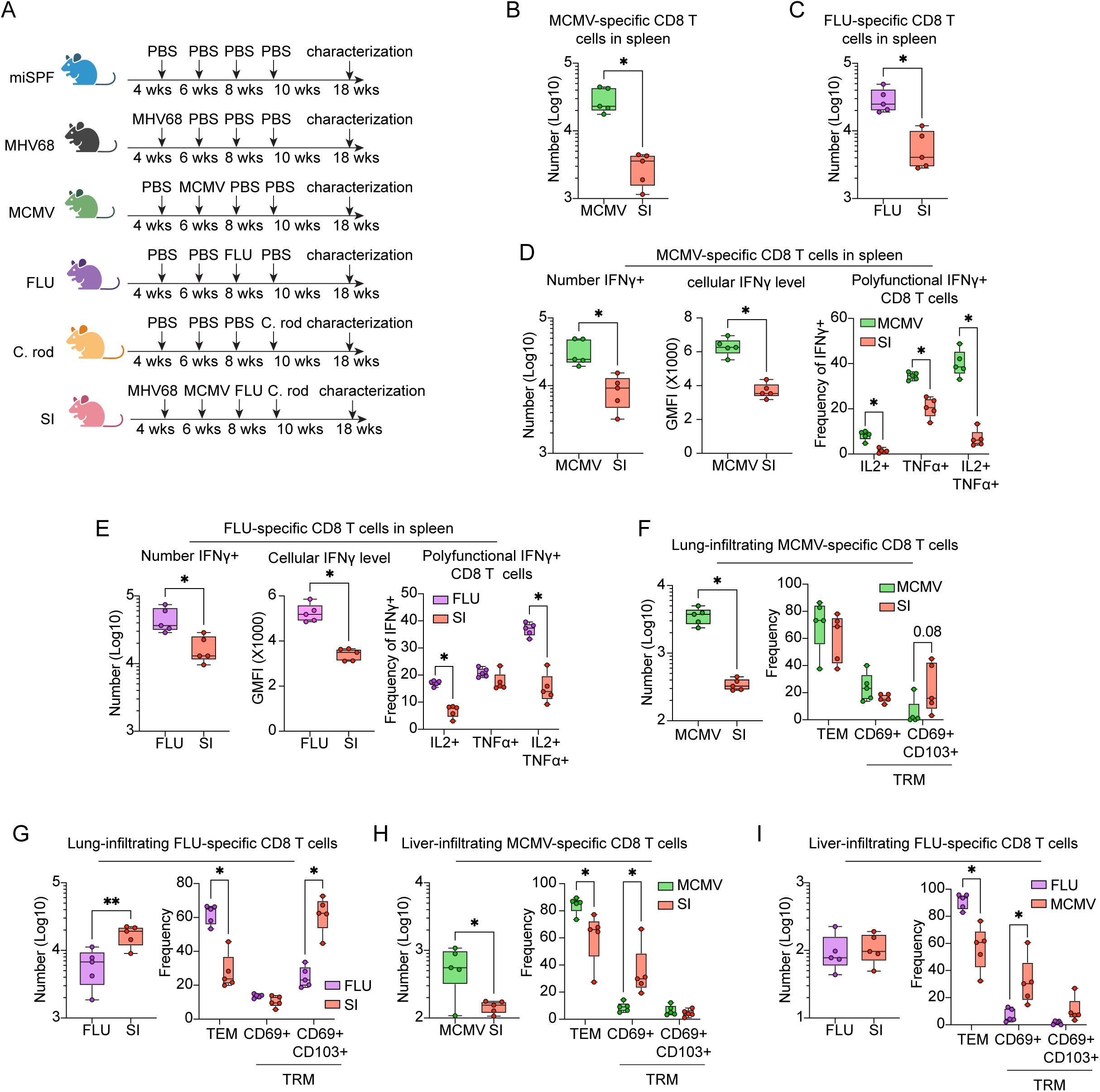
Sequential infection expands total memory while reshaping antigen-specific CD8 T cell responses. **(A)** Experimental design comparing SI with individual pathogen infections. **(B, D)** Splenic MCMV-specific CD8 T cell responses, including cell number, memory subset distribution, and cytokine production following peptide restimulation **(C, E)** Splenic influenza-specific CD8 T cell responses, including cell number, memory subset distribution, and cytokine production following peptide restimulation. **(F–G)** Number and phenotype of lung-infiltrating MCMV-specific and influenza-specific CD8 T cells. **(H–I)** Number and phenotype of liver-infiltrating MCMV-specific and influenza-specific CD8 T cells. Data represent two independent experiments with 4-5 mice per group. Circles represent individual mice. In box-and-whisker plots, boxes indicate the interquartile range, center lines indicate the mean, and whiskers denote the minimum and maximum values. Statistical significance determined by unpaired t-test. *, p<0.05.

Despite this expansion, the magnitude of individual antigen-specific memory populations was reduced. Compared with singly infected mice, SI mice contained fewer MCMV-specific and influenza-specific CD8 T cells in the spleen (Fig. 3B, 3C). This numerical reduction was accompanied by a decrease in the number of IFNγ producing cells and less IFNγ produced per cell accompanied by diminished polyfunctionality, as IFNγ+ cells produced less TNFα, and IL-2 (Fig. 3D, 3E). These findings reveal that successive infections expand total memory while constraining the magnitude and function of individual memory responses.

This phenomenon extended to non-lymphoid tissues but was modified by tissue tropism and infection route. In the lung, the two respiratory infections, MHV68 and influenza, drove substantially greater immune infiltration and memory differentiation than MCMV or *C.rod* (Fig. S4C), highlighting the dominant influence of local infection history on tissue immunity. As observed in the spleen, SI mice exhibited the largest expansion of lung-infiltrating CD8 TEM and TRM populations (Fig. S4C). However, pathogen-specific responses diverged. MCMV-specific CD8 T cells were reduced in SI mice (Fig. 3F), indicating that the diminished splenic response could not be explained by redistribution into peripheral tissues. In contrast, influenza-specific CD8 T cells were increased and preferentially differentiated into CD69⁺CD103⁺ TRM cells (Fig. 3G), suggesting that local antigen encounter can selectively preserve or even enhance tissue-resident memory despite ongoing accumulation of responses to unrelated pathogens. A similar pattern was observed in the liver. Sequential infection increased overall CD8 T cell infiltration (Fig. S4D), while influenza- and MCMV-specific cells adopted liver-appropriate CD69⁺CD103⁻ TRM phenotypes (Fig. 3H, 3I). Thus, successive infections expand total memory while constraining the magnitude and function of individual memory responses.

### Sequential infection reprograms adaptive immune cell states through coordinated inflammatory imprinting

To define how cumulative pathogen exposure reshapes immune cell states, we performed single-cell RNA sequencing (scRNA-seq) on splenocytes from 18-week-old SI and miSPF mice, alongside 7-week-old SPF controls to distinguish infection-driven from age-related changes. Major immune populations were identified (Fig. S5A), and analyses focused on adaptive immune populations, which exhibited the most pronounced transcriptional remodeling.

### CD8 T cells: inflammatory imprinting of naïve cells and effector remodeling of memory

Splenic CD8 T cells were re-clustered into naïve, TCM, and TEM populations, with minor populations (<5%) excluded from further analysis (Fig. 4A). Comparison of 18-week SI and miSPF mice revealed extensive infection-driven remodeling, whereas aging alone induced relatively few transcriptional changes, particularly within memory populations. These findings indicate that cumulative pathogen exposure, rather than age, is the dominant determinant of memory CD8 T cell transcriptional state.

**Figure 4.**
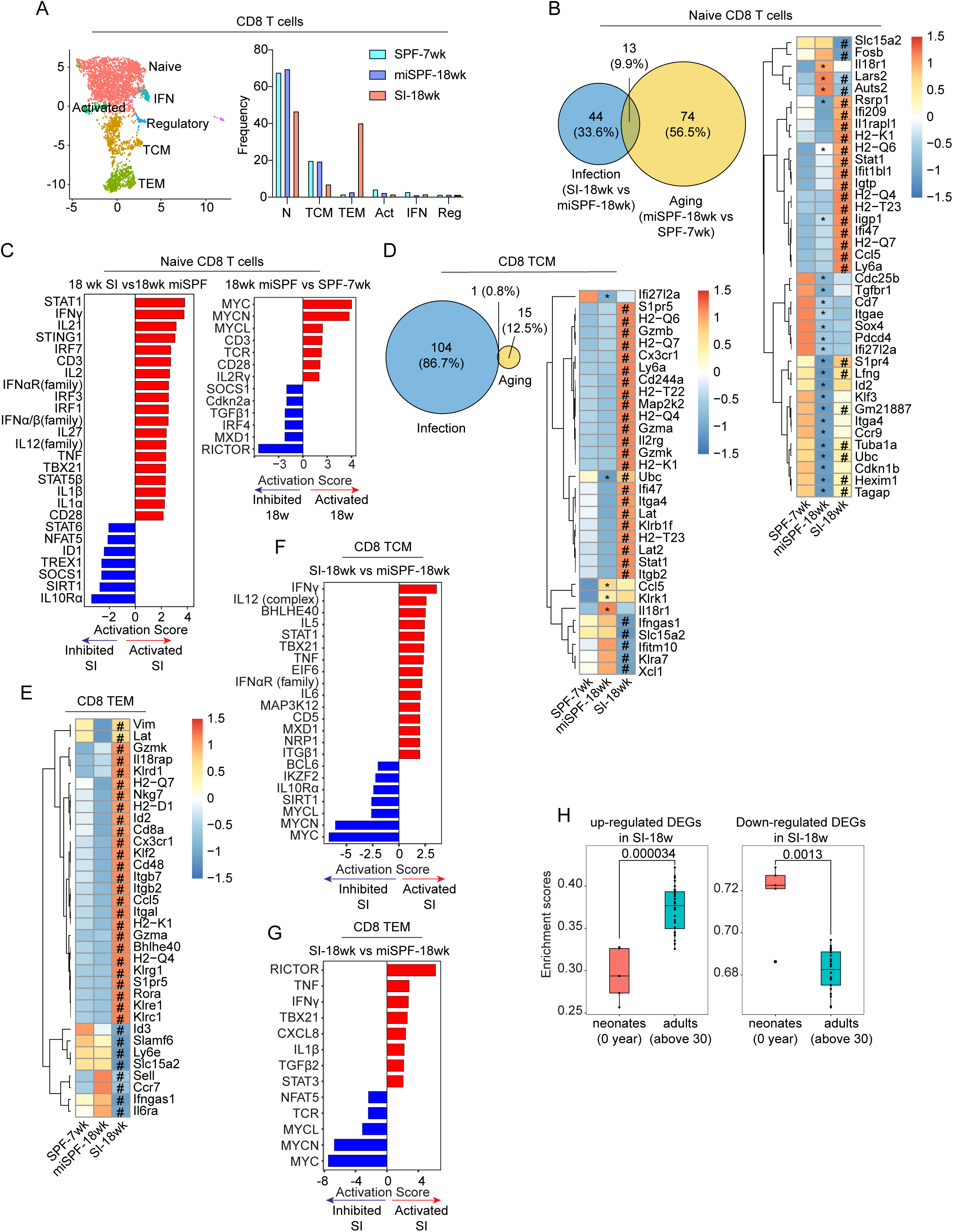
Sequential infection imprints naïve CD8 T cells and promotes effector remodeling of memory CD8 T cells. **(A–G)** scRNA sequencing analysis of splenic CD8 T cells from 18-week-old SI mice, 18-week-old miSPF mice, and 7-week-old SPF controls. **(A)** UMAP visualization of CD8 T cell subsets and their relative frequencies. **(B)** Venn diagrams showing DEGs associated with infection (blue; 18-week SI vs. 18-week miSPF) or aging (yellow; 18-week miSPF vs. 7-week SPF) in naïve CD8 T cells. Heatmap shows scaled expression (Z-score) of selected DEGs. **(C)** Predicted upstream regulators in naïve CD8 T cells identified by IPA. Red indicates predicted activation and blue indicates predicted inhibition. **(D)** Venn diagrams showing infection- and aging-associated DEGs in CD8 TCM cells, with heatmap of selected DEGs. **(E)** Heatmap of selected DEGs in CD8 TEM cells. **(F, G)** Predicted upstream regulators in CD8 **(F)** TCM and **(G)** TEM. **(H)** Expression of human orthologs of SI-associated CD8 TEM DEGs in human CD8 TEM cells across different life stages. Data represent pooled samples from at least four mice per group. *, DEGs for aging comparisons (18-week miSPF vs. 7-week SPF). #, DEGs for infection comparisons (18-week SI vs. 18-week miSPF). Method used for cut-off for DEGs is described in the method session.

Naïve CD8 T cells exhibited substantial transcriptional remodeling following sequential infection. Although aging accounted for a portion of the observed changes, infection and aging regulated largely distinct gene programs with minimal overlap (Fig. 4B). Infection preferentially induced genes associated with migration (*S1pr4, Ccl5)*, interferon signaling (*Ly6a, Stat1, Ifit1bl1, Ifi47, Ifi209, Igtp*), antigen presentation, and immune activation (Fig. 4B). Pathway and upstream regulator analyses further revealed broad activation of inflammatory networks, including IFNα/β, IFNγ, IL-1, IL-12, TNF, STING, and STAT1/IRF signaling, in combination with reduced IL10 signaling (Fig. 4C, S5B), consistent with inflammatory programs previously reported in dirty mice and adult humans (7, 12). In contrast, aging favored pathways linked to proliferation and survival (e.g., MYC, MYCN activation, and TGFβ inhibition), highlighting a fundamental divergence between inflammatory imprinting and age-related homeostasis. Thus, despite retaining a phenotypically naïve appearance, CD8 T cells in SI mice acquire a durable inflammation-experienced transcriptional program, suggesting that cumulative pathogen exposure conditions the naïve repertoire prior to future antigen encounter.

Memory CD8 T cells exhibited minimal aging-associated changes (15 DEGs in TCM and none in TEM) (Fig. 4D, 4E), indicating that their transcriptional state is primarily shaped by infection. In contrast, infections drove extensive reprogramming of both TCM and TEM populations, enriching genes associated with cytotoxicity (KLR and granzyme families), migration (*Cx3cr1, S1pr5, Ccl5, Ccr7, Itgb7*), interferon responsiveness, and antigen presentation, indicating a shift toward enhanced effector function (Fig. 4D, 4E). Interestingly, SI TCM cells upregulated Itga4 (CD49d) (Fig. S5C), a marker that distinguishes infection-induced memory from virtual memory populations (33), demonstrating that SI-generated memory is qualitatively distinct from memory-like populations that arise in SPF mice. Pathway analysis revealed activation of IFNγ, IFNα/β, IL-12, TNF, STAT1, TBX21, and GAIT-associated regulatory networks (Fig. 4F, 4G, S5D). In contrast, miSPF memory cells preferentially expressed biosynthetic, ribosomal, and MYC-driven programs, indicating that cumulative pathogen exposure shifts memory CD8 T cells away from growth-associated states and toward sustained effector function.

To determine whether these transcriptional changes increased similarity to human immunity, we mapped human orthologs of SI-associated CD8 TEM genes and examined their expression across human life stages (34). Genes upregulated in SI mice were significantly enriched in adult human CD8 TEM cells, whereas genes downregulated in SI mice were preferentially expressed in neonatal cells (Fig. 4H). Thus, sequential infection drives coordinated CD8 T cell remodeling in which naïve cells acquire an inflammation-experienced state and memory cells adopt enhanced effector programs that more closely resemble adult human immunity.

### CD4 T cells: inflammatory imprinting and Th1 polarization

Re-clustering of CD4 T cells identified naïve, Th1, Tfh, Tfh expressing high *Itga4* (Tfh_Itga4), Treg, interferon-responsive, and proliferating populations (Fig. 5A). Consistent with our phenotypic analyses, SI mice exhibited increased Th1 cells and reduced naïve CD4 T cells (Fig. 5A). As observed in CD8 T cells, aging contributed relatively little to the overall transcriptional landscape, whereas infection-induced remodeling was concentrated within naïve and Th1 populations (Fig. S6A). Sequential infection induced inflammatory and activation-associated transcriptional programs in both naïve and Th1 cells, including IFNα/β, IL-27, STING, IL-15, and TCR signaling pathways, together with activation of STAT1- and TOX-associated regulatory networks (Fig. 5B, 5C, S6B, S6C). These changes closely mirrored those observed in CD8 T cells, indicating that cumulative pathogen exposure induces a shared inflammatory adaptation program across T cell lineages. In parallel, pathways associated with Th2 and regulatory differentiation, including IL-4, STAT6, and IL-10 signaling, were suppressed (Fig. 5B, S6C), consistent with the expansion of Th1 cells observed at the protein level. Similar to CD8 memory cells, Th1 cells also exhibited reduced MYC-family and translation-associated programs, suggesting a common adaptation toward enhanced effector function and reduced biosynthetic activity. In contrast, aging promoted TCR/CD28 signaling and protein translation while suppressing interferon-associated pathways (Fig. S6B), again highlighting the distinct effects of infection and age. Thus, cumulative pathogen exposure reprograms CD4 and CD8 T cells toward a durable interferon-associated state while reinforcing Th1 differentiation.

**Figure 5.**
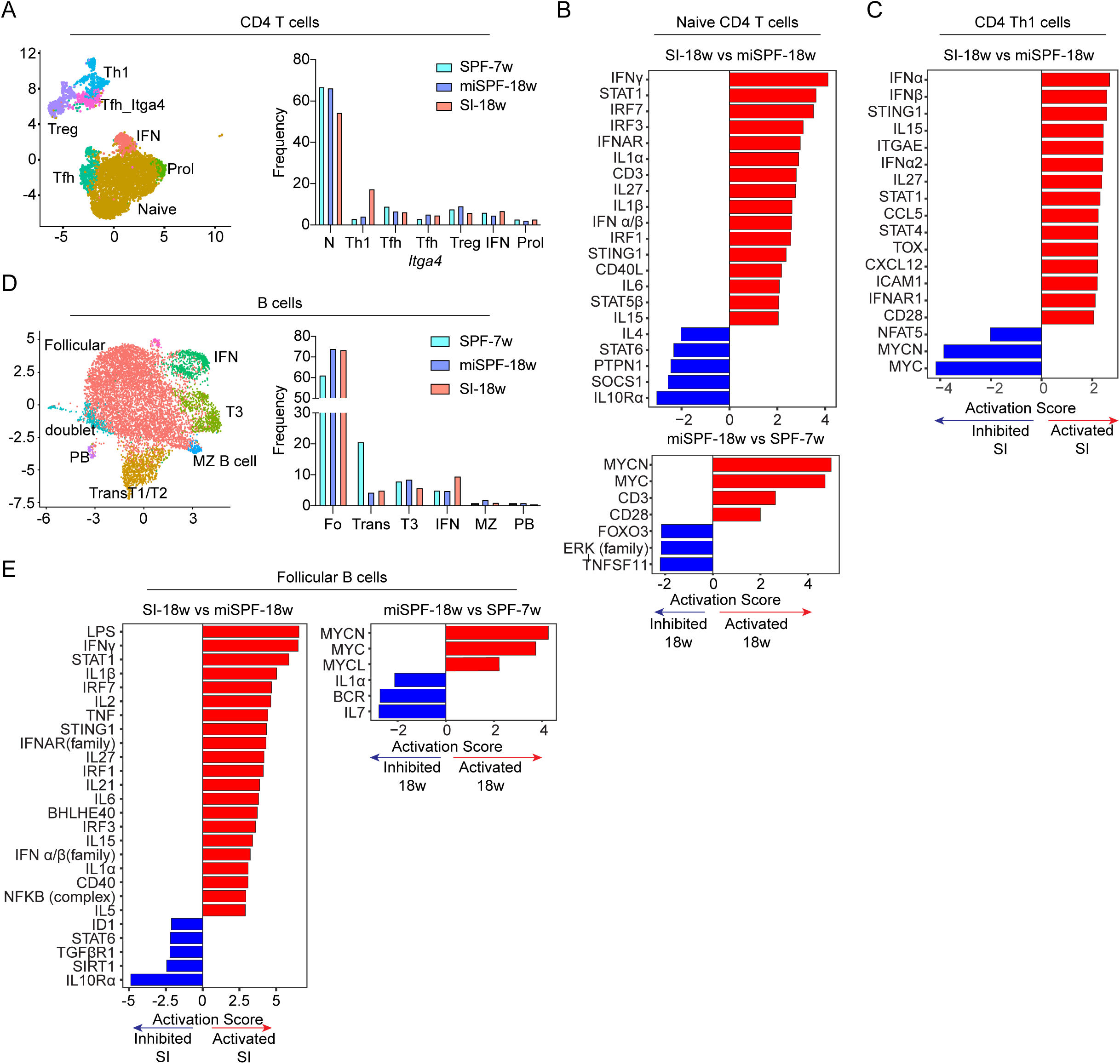
Sequential infection induces coordinated inflammatory reprogramming of CD4 T cells and B cells. **(A–E)** scRNA-seq analysis of CD4 T cells (**A-C**) and B cells (**D, E**) from 18-week-old SI mice, 18-week-old miSPF mice, and 7-week-old SPF mice. **(A)** UMAP visualization and relative frequencies of CD4 T cell subsets. **(B)** Predicted upstream regulators in naïve CD4 T cells associated with infection (left; 18-week SI vs. 18-week miSPF) and aging (right; 18-week miSPF vs. 7-week SPF). Red indicates predicted activation and blue indicates predicted inhibition. **(C)** Predicted upstream regulators in Th1 cells associated with sequential infection. **(D)** UMAP visualization and relative frequencies of B cell subsets. Fo: Follicular B cells; Trans: Transitional T1/T2 B cells; MZ: Marginal Zone B cells; PB: Plasmablasts. **(E)** Predicted upstream regulators in follicular B cells associated with infection and aging. Data represent pooled samples from at least four mice per group.

### B cells: inflammation-experienced reprogramming and enhanced antigen presentation

We next examined B cell populations, identifying follicular (Fo), transitional (T1/T2), T3, marginal zone (MZ), interferon-high (IFN), and plasmablast (PB) subsets (Fig. 5D). Consistent with age-associated maturation, both 18-week SI and miSPF mice exhibited increased follicular B cells and reduced transitional populations relative to 7-week controls (Fig. 5D). However, infection-driven transcriptional remodeling was concentrated within the follicular B cell compartment, whereas other subsets exhibited relatively few differentially expressed genes (Fig. S6D). Follicular B cells from SI mice displayed activation of inflammatory cytokine networks, including IFNγ, IFNα/β, IL-1, IL-6, IL-15, IL-27, and TNF signaling, together with enhanced antigen presentation, BCR signaling, and activation-associated transcriptional programs (Fig. 5E, S6E). These changes were largely absent in aging comparisons, which instead favored cell-cycle progression, ribosomal activity, and protein translation while suppressing inflammatory pathways (Fig. 5E, S6E). Thus, the inflammatory imprint induced by sequential infection extends beyond T cells to the B cell compartment, promoting enhanced antigen presentation and immune activation while reinforcing coordinated remodeling across adaptive immunity. Thus, cumulative pathogen exposure coordinately reprograms adaptive immune cells to establish an inflammation-experienced immune landscape that more closely resembles pathogen-experienced adult humans than conventional SPF mice.

### Modest and transient alterations in the gastrointestinal microbiome following sequential infection

Because infections can influence the gastrointestinal (GI) microbiota, we examined microbial composition in SI and miSPF mice by 16S rRNA sequencing of stool collected at 10 weeks of age (prior to *C. rodentium* infection) and again at 18 weeks. Principal coordinates analysis (PCoA) revealed a modest shift in microbial composition in SI mice at 10 weeks relative to miSPF controls (Fig. S7A), indicating that sequential viral infections and the associated inflammatory responses can influence gut microbial communities even before enteric infection. However, these changes were limited in magnitude.

*C. rodentium* infection did not significantly alter alpha diversity (Fig. S7B), and the dominant bacterial families, including Muribaculaceae and Lactobacillaceae, remained largely unchanged between groups at both time points (Fig. S7C), suggesting preservation of overall community structure. More subtle taxonomic differences were observed longitudinally. Akkermansiaceae was transiently enriched in SI mice at 10 weeks but returned to baseline by 18 weeks (Fig. S7D). Given the established role of *Akkermansia muciniphila* in maintaining intestinal barrier integrity and limiting inflammation (35), this increase may reflect a compensatory response to inflammation induced by the preceding viral infections. In contrast, Acutalibacteraceae was modestly enriched in SI mice at 18 weeks (Fig. S7D), consistent with previous reports linking this family to infection-associated metabolic adaptation (36), suggesting potential shifts in metabolic activity. At the species level, *Muribaculum gordoncarteri* and UBA3263 sp001689615 were reduced in SI mice prior to *C.rod* infection but recovered by 18 weeks (Fig. S7E). In summary, sequential infections induced minor changes on the microbial composition without evidence of broad or sustained dysbiosis. Similarly, reductions in *Muribaculum gordoncarteri* and UBA3263 sp001689615 observed prior to *C. rodentium* infection resolved by 18 weeks (Fig. S7E), indicating that most infection-associated alterations were not sustained. Thus, despite extensive remodeling of immune composition, function, and transcriptional state, sequential infection induced only modest and largely transient alterations in the gastrointestinal microbiota, arguing that the durable immune changes observed in SI mice are unlikely to be driven by broad microbiome disruption.

### SI mice recapitulate functional immune responses of adult humans, and alter the immune response to acute and chronic infections

Having established that sequential infection induces durable phenotypic, transcriptional, and hematopoietic remodeling, we next asked whether these changes alter immune function. We first tested whether SI mice recapitulate a classic example of divergence between SPF mice and adult humans: the response to CD28 super-agonist. Anti-CD28 agonists were originally developed as immunomodulatory therapies because they promote Treg expansion and suppress autoimmunity in SPF mice without significant toxicity (4). However, the catastrophic cytokine storm observed in the TGN1412 clinical trial revealed a profound disconnect between mouse and human immune responses (5). Importantly, environmentally experienced (“dirty”) mice recapitulate the human response to CD28 super-agonist, indicating that immune experience rather than species-specific biology is a key determinant of this divergent outcome (8, 37). Consistent with previous reports, anti-CD28 stimulation in miSPF mice induced robust Treg expansion with minimal systemic inflammation (Fig. 6A, 6B). In contrast, SI mice mounted a rapid TNFα response while exhibiting little Treg expansion (Fig. 6A, 6B), closely resembling responses reported in “dirty” mice and adult humans. Thus, cumulative pathogen exposure confers functional immune maturation, enabling laboratory mice to respond to immunological perturbation in a more human-like manner.

**Figure 6.**
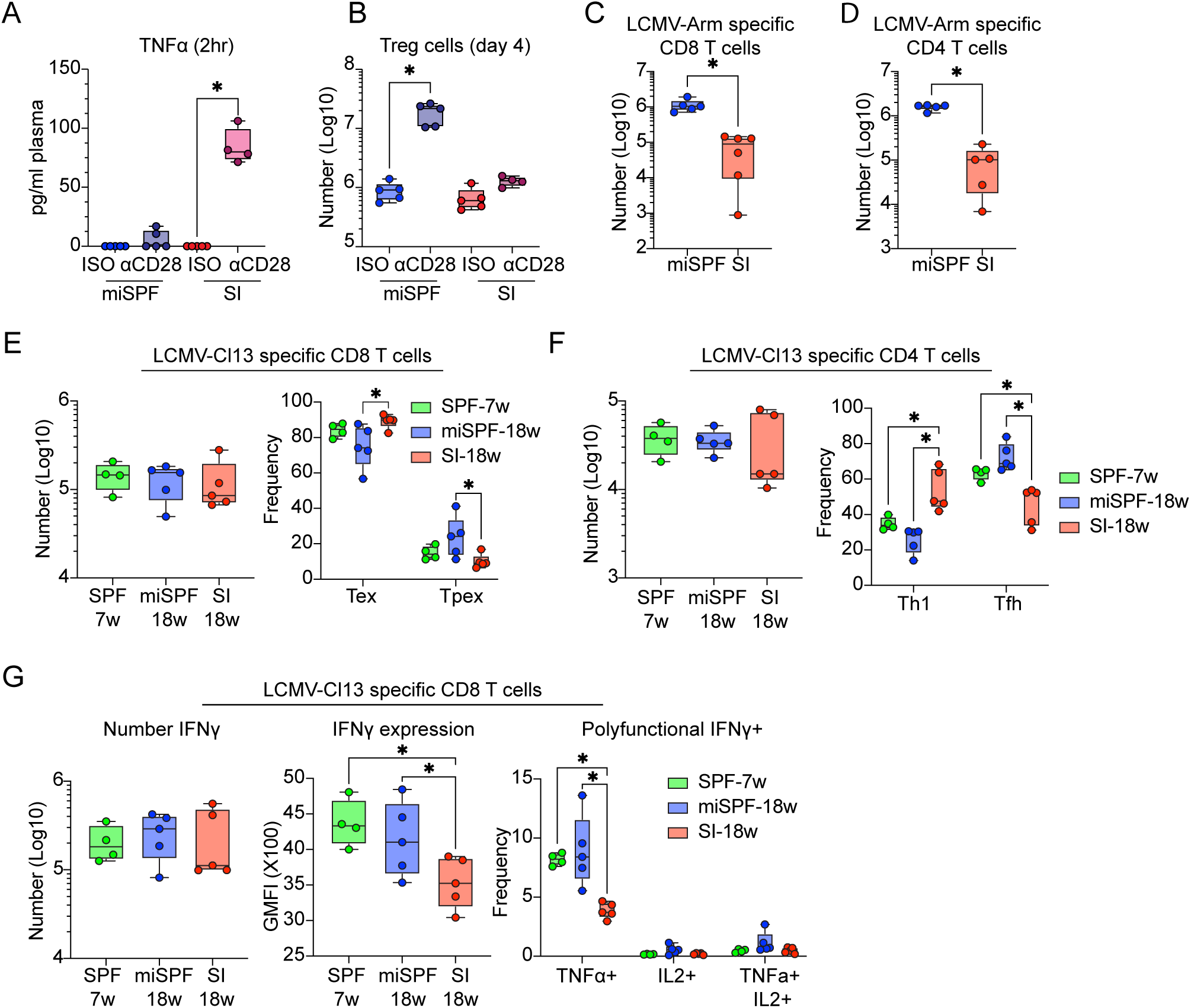
SI recapitulates human-like immune responses and reshapes subsequent antiviral immunity. **(A, B)** Response of 18-week-old miSPF and SI mice to anti-CD28 agonist treatment **(A)** Serum TNFα levels 2 hours after anti-CD28 agonist treatment. **(B)** Number of splenic Treg cells 4 days after anti-CD28 agonist treatment. **(C, D)** Splenic responses to acute LCMV-Armstrong (LCMV-Arm) infection. Quantification of splenic **(C)** LCMV-GP33–41-specific CD8 T cells and **(D)** GP61–80-specific CD4 T cells at day 9 post-infection. **(E-G)** Splenic responses to chronic LCMV-Cl13 infection. At day 9 post-infection, **(E)** number and differentiation of splenic LCMV-GP33–41-specific CD8 T cells, **(F)** number and differentiation of splenic LCMV-GP61–80-specific CD4 T cells, and **(G)** cytokine production by LCMV-GP33–41-specific CD8 T cells following ex vivo peptide restimulation. Data represent two independent experiments (≥4 mice/group), except for the cytokine restimulation in panel G, which was performed only once with at least 4-5 mice per group. Circles represent individual mice. In box-and-whisker plots, boxes indicate the interquartile range, center lines indicate the mean, and whiskers denote the minimum and maximum values. Statistical significance was determined using an unpaired two-tailed t test. *, p<0.05.

We next investigated how the inflammation-experienced immune landscape established by sequential infection influences responses to new viral challenges. To this end, SI and miSPF mice were infected with LCMV-Armstrong (LCMV-Arm) or LCMV-Clone 13 (LCMV-Cl13), to generate an acute or chronic viral infection, respectively (38). During acute LCMV-Arm infection, SI mice generated fewer LCMV-specific CD8 T cells than miSPF controls (Fig. 6C). This reduction occurred without major changes in SLEC versus MPEC differentiation (Fig. S8A), indicating that sequential infection primarily affected the magnitude rather than the developmental trajectory of the response to acute infection. Consistent with reduced expansion, IFNγ production was reduced both in the total number of producing cells and on a per-cell basis, although cytokine polyfunctionality was largely preserved (Fig. S8B). Likewise, LCMV-specific CD4 T cell responses were diminished in SI mice, while helper T cell differentiation remained largely unchanged (Fig. 6D, S8C, S8D). These findings mirror our earlier observations that sequential infection expands total memory while limiting individual antigen-specific responses, suggesting that cumulative pathogen exposure constrains the magnitude of *de novo* T cell responses despite establishing a larger overall memory compartment.

In contrast, during chronic LCMV-Cl13 infection, sequential infection primarily altered T cell differentiation rather than expansion. Interestingly, both miSPF and SI mice longitudinally controlled virus more effectively than 7-week-old SPF controls (Fig. S8E), indicating the impact of age on control of chronic viral infection. At day 9 post-infection, SI and miSPF mice generated comparable numbers of virus-specific CD8 and CD4 T cells (Fig. 6E, 6F), indicating that initial priming was preserved. However, virus-specific CD8 T cells in SI mice preferentially differentiated into terminally exhausted (Tex) cells at the expense of progenitor exhausted (Tpex) cells and exhibited reduced cytokine production (Fig. 6E, 6G), suggesting increased exhaustion in SI mice. Similarly, virus-specific CD4 T cells displayed enhanced Th1 differentiation and reduced Tfh development (Fig. 6F). These shifts closely parallel the broader immune remodeling observed throughout the study, including increased Th1 polarization, enhanced inflammatory imprinting, and reduced support for Tfh programs. Thus, rather than altering the magnitude of the chronic antiviral response, sequential infection redirects its differentiation trajectory toward terminal effector and exhausted states. Thus, cumulative pathogen exposure establishes an inflammation-experienced immune landscape that reshapes both the magnitude and differentiation of subsequent antiviral responses, linking immune history to future immunity.

## DISCUSSION

Specific pathogen-free mice have been indispensable for defining fundamental principles of immunology, yet their limited antigenic experience remains a major barrier to modeling adult human immunity. While previous studies established that SI promotes memory T cell accumulation and broad immune maturation (12, 39), how cumulative pathogen exposure reshapes immune homeostasis and future immune responses has remained poorly understood. Here, we show that SI drives coordinated remodeling across the immune system, extending from hematopoietic progenitors to innate immune populations, adaptive lymphocytes, and tissue-resident immunity. Collectively, these changes establish a durable inflammation-experienced immune landscape that more closely resembles pathogen-experienced adult humans than conventional SPF mice.

A central observation of this study is that cumulative pathogen exposure does not simply increase immune memory but fundamentally alters its organization and maintenance. SI progressively expanded the total memory T cell compartment while reducing the magnitude and functionality of individual antigen-specific responses, demonstrating that immune experience is not merely additive. These effects were particularly evident in non-lymphoid tissues, where sequential infection promoted the accumulation of tissue-resident memory T cells and established mature immune landscapes within the lung and liver. Because tissue residency is associated with enhanced persistence and occupation of specialized survival niches (40, 41), repeated inflammatory exposure may progressively redistribute memory populations from circulation into long-lived resident compartments. Such adaptations may more closely reflect human immunity, where lifelong pathogen exposure continuously reshapes immune composition and function.

Beyond mature immune populations, SI altered the cellular foundation from which future immune responses are generated. SI remodeled hematopoietic stem and progenitor compartments, selectively reducing myeloid-biased progenitors while preserving lymphoid progenitors, and induced coordinated transcriptional reprogramming across adaptive immune cells. Most notably, naïve CD4 and CD8 T cells acquired interferon-associated transcriptional programs despite retaining a phenotypically resting state, while memory T cells adopted enhanced effector programs and follicular B cells acquired signatures of inflammatory activation and antigen presentation. These adaptations were largely distinct from age-associated changes and more closely resembled immune programs observed in pathogen-experienced adult humans, suggesting that immune history not only shapes pre-existing immunity but also conditions how future immune responses develop following new antigen encounter.

The functional consequences of this remodeling became apparent following immunological challenge. Consistent with previous observations in dirty mice (8), SI mice responded to CD28 super-agonist with inflammatory cytokine production rather than Treg expansion, recapitulating a hallmark difference between SPF mice and adult humans. Likewise, SI altered responses to both acute and chronic viral infection. During acute LCMV infection, virus-specific CD4 and CD8 T cell expansion was reduced despite preserved differentiation, whereas during chronic infection, T cell differentiation shifted toward terminal effector and exhausted states at the expense of progenitor populations. These findings mirror the transcriptional and phenotypic changes observed at baseline and support a model in which inflammatory imprinting of naïve lymphocytes influences subsequent cell fate decisions.

Although SI mice share many features with other environmentally exposed or dirty mouse models, important differences highlight how the nature of pathogen exposure shapes immune outcomes. In contrast to co-housed and rewilded mice, SI mice exhibited reduced circulating granulocytes and monocytes, likely reflecting differences in pathogen composition and repeated inflammatory signaling that can suppress hematopoietic output, including through IFN-I–dependent mechanisms (14, 16, 42, 43). Thus, immune experience is not a single biological state but rather a continuum influenced by the identity, sequence, timing, and persistence of prior infections.

These findings have important implications for human immunity and translational research. Elevated baseline inflammatory signaling, extensive tissue-resident memory populations, and altered cytokine responsiveness are hallmarks of pathogen-experienced adult humans and are increasingly recognized as determinants of vaccine efficacy, infection outcomes, autoimmunity, and cancer immunotherapy (44–48). By recapitulating many of these features, SI provides a controlled and reproducible platform for investigating how infection history shapes immune homeostasis, disease susceptibility, and therapeutic responsiveness. Ultimately, cumulative pathogen exposure reprograms hematopoiesis, immune cell state, and the differentiation of future immune responses, establishing immune history as a fundamental determinant of immune function.

## AUTHOR CONTRIBUTIONS

M.G., D.G.B. designed research. M.G., Y.H.B., H.J.E., W.X., M.Y.R.L., and J.E.M.O. performed experiments. T.A.R. provided viruses and protocols. A.M. provided *C. rodentium* and protocols. F.N.I. and L.J.E. helped with Spectral Flow panel design and acquisition. M.T.C., M.B. and Z.Q.L performed 16s rRNA sequencing. M.G., Y.H.B., D.A-R., R.Q and D.G.B analyzed data. M.Y.R.L., W.X., T.L.M. and D.G.B. contributed experimental direction, insight, and discussion. M.G. and D.G.B. wrote the manuscript.

## ACKNOWLEDGEMENTS

We thank past and present members of the Brooks and McGaha laboratories for technical help and discussion. This work was supported by the Canadian Institutes of Health Research (CIHR) Grants FDN148386, PJT191843 (D.G.B), the National Institutes of Health (NIH) grant R21 CA296062 (D.G.B and T.A.R), and the Scotiabank Research Chair to D.G.B.

## METHODS

### Lead contact

Further information and requests for resources and reagents should be directed to and will be fulfilled by the Lead Contact, David Brooks.

## Materials Availability

Requests for resources and reagents should be directed to David Brooks.

### Mice

C57BL/6 mice were purchased from Jackson Laboratories or the rodent breeding colony at the Princess Margaret Cancer Centre. All mice were housed under specific-pathogen free conditions. Mouse handling conformed to the experimental protocols approved by the Animal Care Committee at the Princess Margaret Cancer Center / University Health Network. Unless otherwise specified, female mice were used at 18 weeks.

### Viral and bacterial infections

Pathogen information and infection strategy are listed in the material and reagent section (Table S1) and Figure 1A. To set up sequentially infected mice: at 4 weeks, SPF mice were infected with 10^5^ plaque forming units (PFU) of MHV68 (strain WUMS) intranasally (in 20ul PBS); at 6 weeks mice, were infected intraperitoneally with 10^5^ PFU (in 100ul PBS) of MCMV (strain SMITH); at 8 weeks, mice were infected intranasally with 10^3^ PFU influenza (H1N1, strain A/Puerto Rico/8/1934) (in 20ul PBS); at 10 weeks, mice were infected by oral gavage with 2x10^9^ colony forming unit (CFU) of *Citrobacter rodentium* (strain DBS100, in 150 ul PBS). Mice were then rested in the same facility for another 8 or 12 weeks before use, while supportive nutrients (ensure) were provided daily after 1 week of *C. rodentium* gavage for 2 weeks. In parallel, miSPF mice were mock infected at the same time points and via the same route with PBS. For LCMV infection, mice were infected intraperitoneally with 2x10^4^ PFU of LCMV-Armstrong or intravenously with 2x10^6^ PFU of LCMV-Clone13.

### In vivo intravascular antibody labelling

For in vivo antibody labeling, pan-CD45 antibody was injected intravenously at 4ug per mouse in 150ul PBS and sacrificed 4 minutes later as described (25).

### Tissue processing

Spleens were harvested and manually dissociated into single cell suspension. Cells were passed through 100-micron filter and then subjected to red blood cell (RBC) lysis at room temperature (RT) for 3 min. Lymph nodes were harvested, manually dissociated into small pieces, and digested in 500ul digestion buffer (RPMI 1640 medium with 10% FBS, 1% HEPES, 1 mg/ml Collagenase IV from Clostridium histolyticum and 0.15 mg/ml DNase I) at 37° for 30 min. Lymph node single cell suspensions were obtained by filtering through the strainer snap cap (Fisher scientific Cat#08-771-23).

Lungs were harvested, manually dissociated into small pieces, and digested in 5ml digestion buffer at 37°C for 1 hour. Lung single cell suspensions were obtained through 100-micron filters and RBC lysis at RT for 4 min. Livers were harvested and manually dissociated into suspension with a tissue smasher. The suspension was passed through 100-micron filter and spin for 15 min with 35% Percoll gradient at 4 °C. Cell pellets were subjected to RBC lysis at RT for 4 min. Bone marrow was harvested from tibia and femur, passed through a 100-micron filter and then subjected to RBC lysis at room temperature for 3 min. To obtain single cell suspension from the brain, perfused tissue was manually processed through 100-micron strainer, and lymphocytes were isolated using Percoll gradient. For all tissues, after RBC lysis cell counts were obtained before proceeding to flow cytometry or CyTOF staining.

### *Ex vivo* peptide stimulation

1-2x10^6^ splenocytes were stimulated with 1µg/mL of MHC class I-restricted peptides in 96-well U-bottom plates. The stimulation was performed in the presence of 50 U/mL recombinant murine IL-2 and 1 µg/mL brefeldin A for 5 hours at 37°C.

### Flow cytometry staining

Single cell suspensions were stained *ex vivo* using antibodies listed in the reagent list. Live/dead staining was done using zombie aqua or zombie NIR. Staining was performed as directed using the Foxp3 Transcription Factor Staining kit. Samples were run on a FACS Lyrics (BD Biosciences) and data analyzed using FlowJo software (BD biosciences). For spectral flow analysis, samples were run on the Cytek Aurora. Cell types in flow cytometry experiments were defined/gated as following. We include this information to promote more consistent definition and improve the reproducibility of data across studies (49).

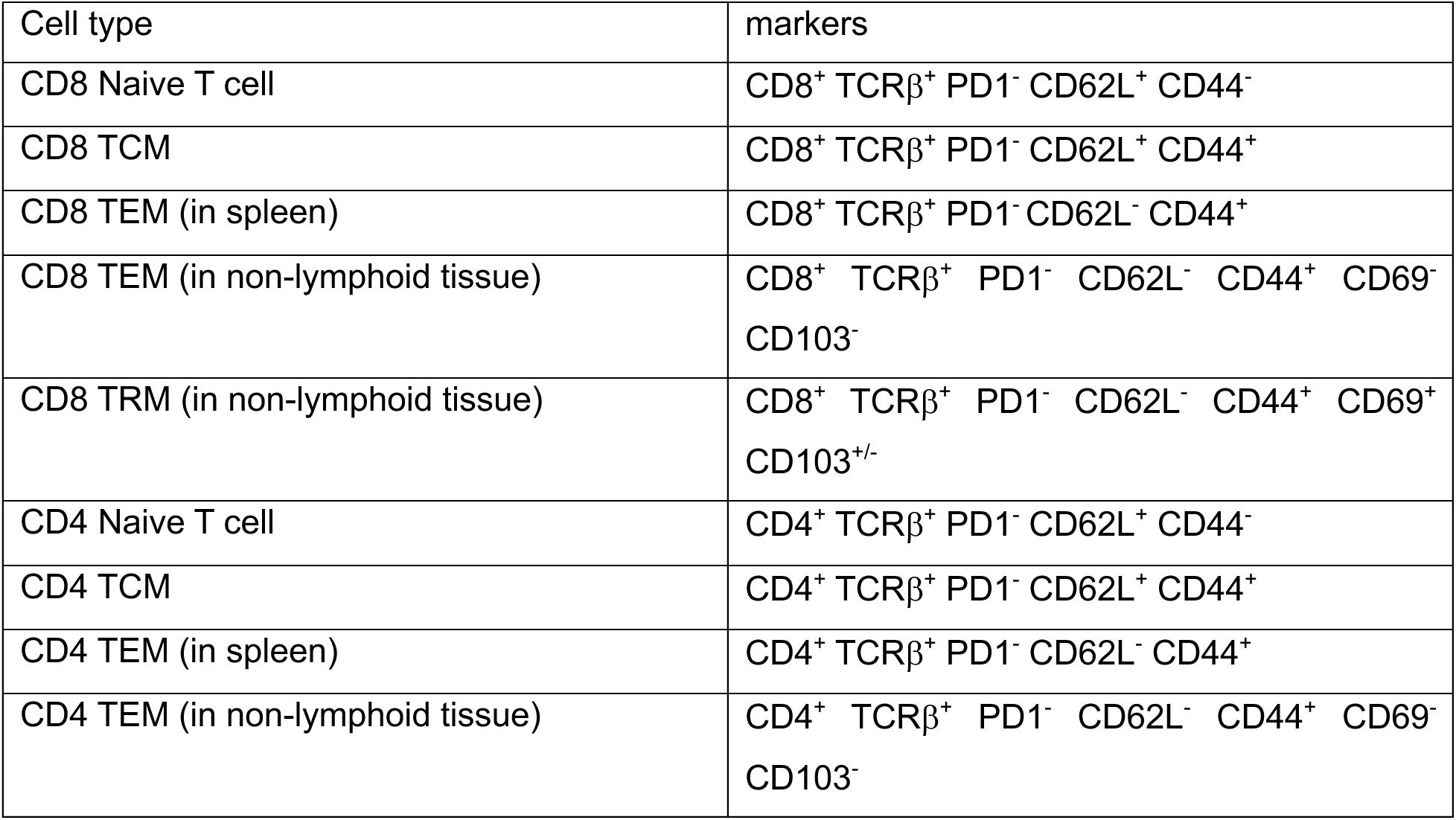
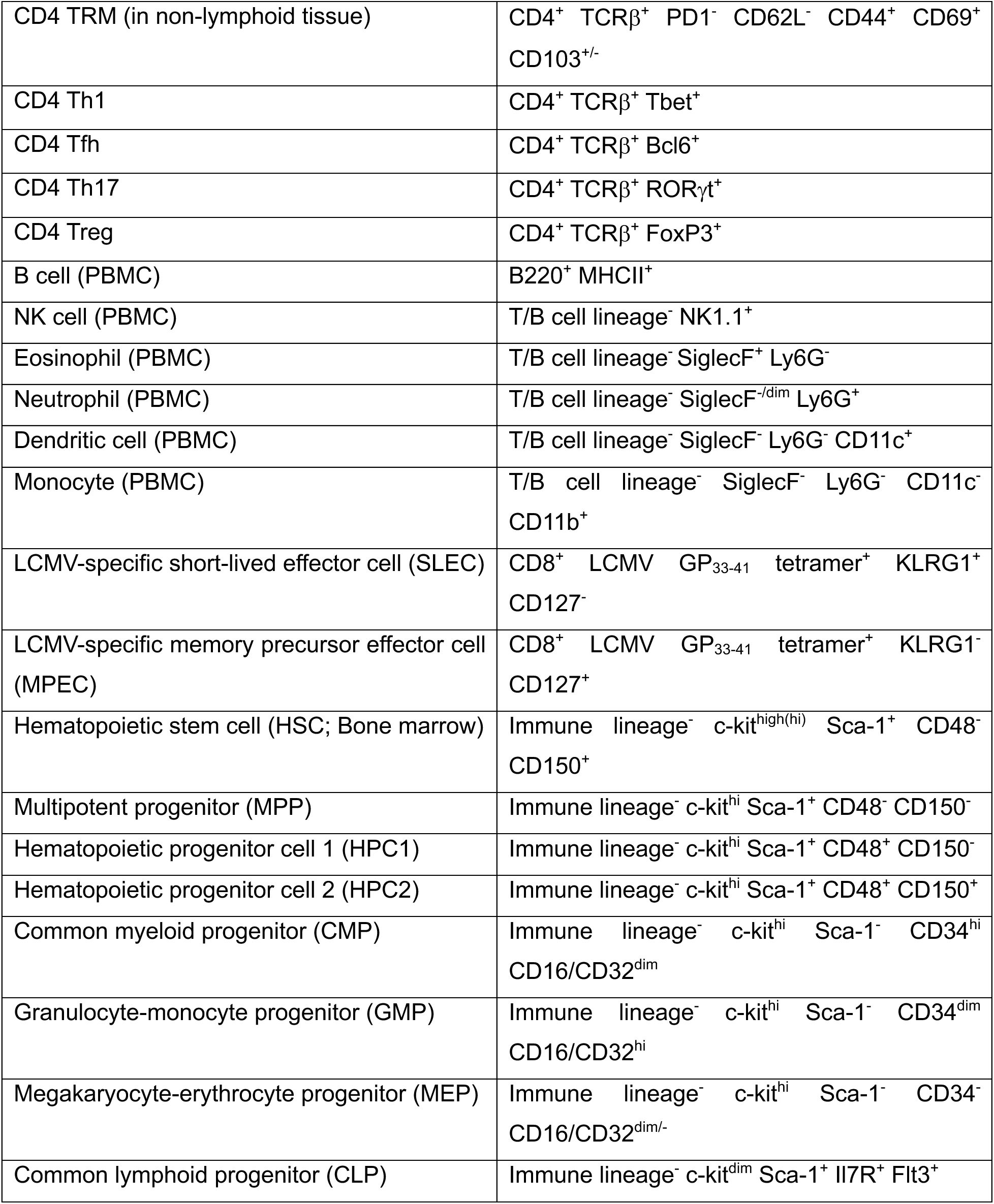

### Mouse total IgG and TNFα ELISA

TNFα ELISA was performed according to Biolgend’s manual. For plasma IgG titers, plates were coated overnight with 2 μg/ml goat anti-mouse IgG. Plates were washed and blocked with 3% BSA. Mouse IgG standard or plasma samples were added and incubated at RT for 2 hours. Then goat anti-mouse IgG HRP (1mg/ml) was added, followed by substrate. The reaction was stopped with 2N H_2_SO_4_. Results were read on ELISA reader at 492 nm.

### Time-of-flight mass cytometry (CyTOF)

CyTOF antibodies are listed in the reagent list. Where indicated, already conjugated antibodies were purchased from Fluidigm. Conjugation of purified antibodies was performed at the SickKids-UHN Flow and Mass Cytometry Facility using the MaxPar Antibody Labeling Kit (Fluidigm). All antibodies were titrated prior to the experiments. During staining, samples were washed with PBS and stained with 12.5μM cisplatin in PBS for 1 min at RT, and quenched with CyTOF staining media (Mg+/Ca+ HBSS containing 2% FBS, 10mM HEPES) and FBS. For antigens sensitive to fixation, cells were FcR blocked and stained with metal tagged antibodies at RT for 10 min. Then, cells were fixed with the Foxp3 Transcription Factor Staining kit and barcoded with Cell-ID 20-Plex Pd Barcoding Kit. Then, combined samples were stained with metal tagged antibodies. Surface antibodies were diluted in CyTOF staining media and cells were stained at 4°C for 30 min. Intracellular antibodies were diluted in the permeabilization buffer from Foxp3 Transcription Factor Staining kit and cells were stained at RT for 30 min. For the DNA stain, cells were incubated overnight in PBS with 0.3% (w/v) saponin, 1.6% (v/v) paraformaldehyde and 1nM iridium. Samples were acquired on a Helios mass cytometer (Fluidigm) at the Princess Margaret Cancer Centre. EQ Four Element Calibration Beads (Fluidigm) were used to normalize signal intensity over time on CyTOF software version 6.7.

### Bioinformatic analyses (CyTOF)

Preprocessing of files was performed using FlowJo Software (v10) software. Samples were manually debarcoded and exported as separated FCS files. Cells were filtered by gating on DNA, singlets, live cells, and raw signal events were then exported as matrices in csv format. Data was then analyzed in R (v 4.1.0). All events were included in dimensionality reduction. Protein expression values were arcsinh transformed using a custom co-factor for each protein before clustering. Phenograph and UMAP were performed using the R implementation of the “Rphenograph” package (v 0.99.1) by JinmiaoChen lab on github (50) and package “umap” (v 0.2.7.0). Differential state and abundance analyses were performed by “diffcyt” package (51) (v 1.14.0).

### Single cell RNA-sequencing

scRNAseq was performed by The Princess Margaret Genomics Centre (PMGC). *Sample preparation and library generation:* Spleens were harvested, processed into single cell suspension, and pooled within each condition. Spleens were pooled from 4-5 mice per group. Samples were prepared as outlined by 10x genomics CG000331_Chromium_Next_GEM_Single_Cell_5_v2_UserGuide and loaded onto 10X single cell chip K. After droplet generation, samples were transferred onto a pre-chilled 96 well plate (Eppendorf), heat sealed and incubated overnight in a Veriti 96-well thermo cycler (Thermo Fisher). The next day, sample cDNA was recovered using Recovery Agent provided by 10x and subsequently cleaned up using a Silane DynaBead (Thermo Fisher) mix as outlined by the user guide. Purified cDNA was amplified for 13 cycles before being cleaned up using SPRIselect beads (Beckman). 5’ v2 cDNA libraries were prepared as outlined by the Single Cell 5’v2 Reagent Kits user guide with modifications to the PCR cycles based on the calculated cDNA input.

- Chromium Next GEM Single Cell 5’ Kit v2, 16 rxns PN-1000263
- Chromium Next GEM Chip K Single Cell Kit, 48 rxns PN-1000286
- Dual Index Kit TT Set A, 96 rxns PN-1000215

*Sequencing*: Library size was determined using Agilent High Sensitivity DNA Kit on the Bioanalyzer instrument (Agilent Technologies). The samples were then quantified using the Qubit dsDNA HS Assay Kit on the Qubit 2.0 Fluorometer (Invitrogen), and subsequently normalized to 1.5 nM using Low TE buffer (Invitrogen). The 1.5 nm pool was denatured using 0.2N NaOH according to Illumina guidelines for 8 minutes at room temperature. Library pool was further diluted to 350 pM using 400mM Tris-HCl. This multiplexed pool was sequenced with the following parameters with an S2 flow cell on the NovaSeq X platform (Illumina): Read 1 – 28 cycles, Read 2 – 91 cycles, Index 1 – 8 cycles. Approximately 10^6^ reads were acquired per cell.

### Bioinformatic Analyses (scRNA-Seq)

Single-cell data was analysed in R (v 4.4.2) and the R Seurat (v 5.3.0) package. *Reads alignment and quantification:* Base calling was performed using Illumina RTA (v 4.29.2) and BCL files were converted to Fastq files using the Illumina bcl2fastq2 (v 4.1.7). Sequencing reads were aligned to the mouse reference genome mm10 (10x Genomics reference bundle refdata-gex-mm10-2020-A) using Cell Ranger (v9.0.1), and gene expression matrices were generated using the Cell Ranger count pipeline.

*Quality control*: Cells considered dead or of poor quality were identified as outliers based on the Median Absolute Deviation (MAD) using the isOutlier function from the scuttle package (v1.16.0). Specifically, dead cells were defined as outliers based on mitochondrial gene percentage (settings: nmads=7, type=”higher”, min.diff=15). Poor-quality cells were identified as outliers based on both number of features and ribosomal gene percentage (nmads=7 and 6 respectively, type=”lower”, log=T). Furthermore, cells were excluded based on the additional following criteria: hemoglobin percentage < 0.1, and log10GenesPerUMI > 0.8 *Doublet identification*: Single-cell data was analysed in R (v 4.4.2) and the R Seurat (v 5.3.0) package (52). Initial clustering for each sample was performed following the standard Seurat workflow (i.e. NormalizeData, FindVariableFeatures, ScaleData, RunPCA, FindNeighbors, and FindClusters) with default parameters and a resolution of 1. Doublet scores were computed by the R package DoubletFinder (v2.0.6) using its standard workflow and default parameters (53). Doublet scores were standardized to Z-scores; cells with Z-score > 2 were flagged as potential doublets. Clusters containing > 10% potential doublets were identified as “doublet clusters” and were each subjected to re-clustering. To identify sub-clusters that are enriched for doublets, we calculated the delta between individual cell Z-scores and the sample median Z-score. Sub-clusters with a modified Z-score > 4 (derived from the median of these deltas) were classified as doublets and excluded from downstream analyses.

The number of cells remaining in each sample following these filters is as follows, and genes expressed in less than 10 cells were excluded from further analyses:

- SPF_7w: 6635 cells
- SPF_18w: 8023 cells
- SI_18w: 6444 cells

*Sample integration*: Individual samples were preprocessed following the standard Seurat workflow (i.e. NormalizeData, FindVariableFeatures, ScaleData, RunPCA). Ribosomal and cell cycle genes were excluded from the set of variable features prior to RunPCA to prevent these genes from influencing downstream clustering results. Samples within each group were integrated using the IntegrateLayers function with the RPCA algorithm (RPCA vignette: https://satijalab.org/seurat/articles/integration_rpca.html?utm_source=chatgpt.com). Nearest-neighbors graph construction was performed using the first principal components that explain 90% of the data variability from the RPCA-integrated reduction. Clustering using the Louvain algorithm and resolution 0.5 identified the main cell types: B cells, CD4 T cells, CD8 T cells, and innate immune cells. Each sub-population was re-integrated and re-clustered using the same workflow to identify fine sub-population using the following resolutions: 0.2, 0.4, 0.2 and 0.2, respectively.

*Differential gene expression*: Differential gene expression analysis was conducted using the Wilcoxon rank-sum test implemented in Seurat, with Bonferroni correction applied for multiple hypothesis testing across predefined cell populations. Genes were considered significantly differentially expressed, if they exhibited an absolute log2 fold change > 0.25 and were expressed in at least 10% of cells in one of the compared groups.

*Human Mouse similarity analysis:* Human genes were mapped to their corresponding mouse orthologs via the babelgene R package (v22.9). Up- and down-regulated DEGs from the mouse SI_18w vs. SPF_18w comparison in CD8 TEM were used to define two functional gene sets. Cellular enrichment scores were quantified using the AddModuleScore_UCell function using the UCell R package (v1.28.0). Statistical significance of UCell scores between human infant (age = 0 years) and adult cohorts (age ≥ 30 years) was evaluated using a two-tailed Wilcoxon rank-sum test for CD8_TEM_CMC1 cells in Human.

*QIAGEN Ingenuity Pathway Analysis (IPA) and Upstream Regulator analysis:* Gene fold change and q-value were used as input for QIAGEN IPA. Analysis was based on genes with a q-value less than or equal to 0.05. The algorithms developed for use in IPA were described by QIAGEN (54).

### 16s ribosomal RNA sequencing and analyses

*Sample collection and processin*g: Stool samples were collected directly into 1.5 ml sterile eppendorf tubes, snap frozen with liquid nitrogen, and kept at -80 °C until use (no fasting or pre-treatment of mice was required). Microbial DNA was purified with ZymoBIOMICS DNA Miniprep Kit (Zymo Research). Amplicon libraries for the V3 and V4 hypervariable regions of the 16S rRNA gene were generated as previously described the Illumina protocol #15044223 Rev. B. Library traces were obtained using a Bioanalyzer 2010 with the High Sensitivity DNA kit (Agilent), and concentrations were measured using the Qubit High Sensitivity DNA kit (ThermoFisher Scientific). Libraries were pooled and diluted to 60pM, with a 5% PhiX (Illumina) spike-in for loading. Sequencing was performed using paired-end 150 bp reads, generating a total of 5 million reads on a MiSeq i100 sequencer (Illumina) at the Keenan Research Centre for Biomedical Sciences (Toronto).

*Data analysis:* Raw paired-end 16S rRNA gene sequencing data were processed using QIIME 2 (v2024.10.1). Reads were imported using the *PairedEndFastqManifestPhred33V2* format and the sequence quality was assessed using demux. Sequences were quality-filtered to a trim length of 150vp and amplicon sequence variants were called from denoised data using Deblur. Phylogenetic trees were constructed using MAFFT and aligned using the FastTree algorithm. Alpha and beta diversity metrics were calculated using a rarefaction depth of 5,000 reads per sample. Alpha diversity was measured using Faith’s Phylogenetic Diversity and unweighted UniFrac distances. Group differences were assessed using Faith’s phylogenetic diversity and the Bray-Curtis distance was used for a Principal Coordinate Analysis. Taxonomic classification was performed using a Naive Bayes classifier trained on the Greengenes (2022.10 release). Feature tables were collapsed at family (level 5), genus (level 6) and species (level 7). These feature tables were tested for differential abundances using ANCOMBC2 (v2.12.0), and taxonomic composition was visualized using bar plots.

### Statistical analyses (other than CyTOF, microbiome, and scRNAseq)

Statistical tests were calculated by GraphPad Prism 9 (GraphPad Software, Inc.) as indicated in the Figure legends.

---

**Table S1.** Pathogens used for sequential infections.

| Pathogen | Strain | Infection method | Relevance to Human |
| --- | --- | --- | --- |
| Murine gammaherpesvirus 68 (MHV68) | WUMS | intranasal | Murine model for the human gammaherpesviruses Epstein-Barr virus (EBV) and Kaposi's sarcoma-associated herpesvirus (KSHV) |
| Murine cytomegalovirus (MCMV) | SMITH | intraperitoneal | Murine model for human cytomegalovirus |
| Influenza A virus (H1N1) | A/PR/8/34 | intranasal | Mouse-adapted influenza A virus |
| *Citrobacter rodentium* (C. rod) | DBS100 | oral gavage | Murine model for *E. coli* infection |

**Figure S1.**
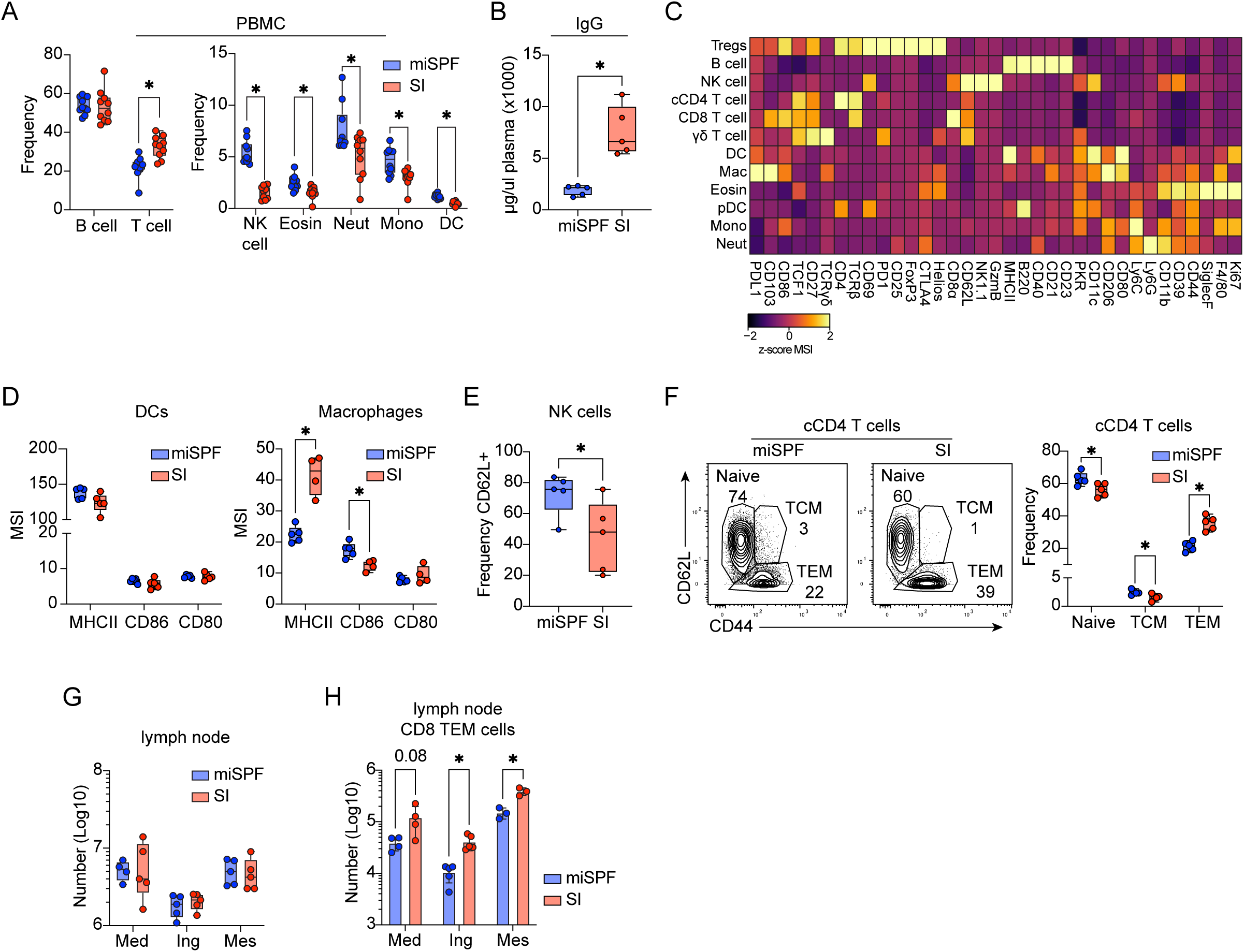
Sequential infection alters immune composition in blood, spleen, and lymph nodes. All analyses performed in 18-week-old miSPF and SI mice. **(A)** Frequency of immune subsets among CD45⁺ peripheral blood mononuclear cells (PBMC) measured by flow cytometry. **(B)** Plasma IgG levels quantified by ELISA. **(C)** Median signal intensity (MSI) of phenotypic markers across splenic immune subsets identified by Phenograph clustering. **(D)** MSI of MHC II, CD80, and CD86 expression on splenic dendritic cells and macrophages. **(E)** Frequency of CD62L expression on splenic NK cells. **(F)** Representative gating and quantification of naïve, TCM, and TEM subsets within FoxP3⁻ conventional (c)CD4 T cells. **(G)** Total number of CD45⁺ immune cells across mediastinal, inguinal, and mesenteric lymph nodes. **(H)** Numerical quantification of CD8 TEM cells in lymph nodes. Data represent at least two independent experiments (≥4 mice/group). Circles in the box and whisker plots represent individual mice. Error bars in whisker plots indicate the highest and lowest value, and the line in the box indicates the mean. Statistical significance determined by unpaired t-test. *, p<0.05.

**Figure S2.**
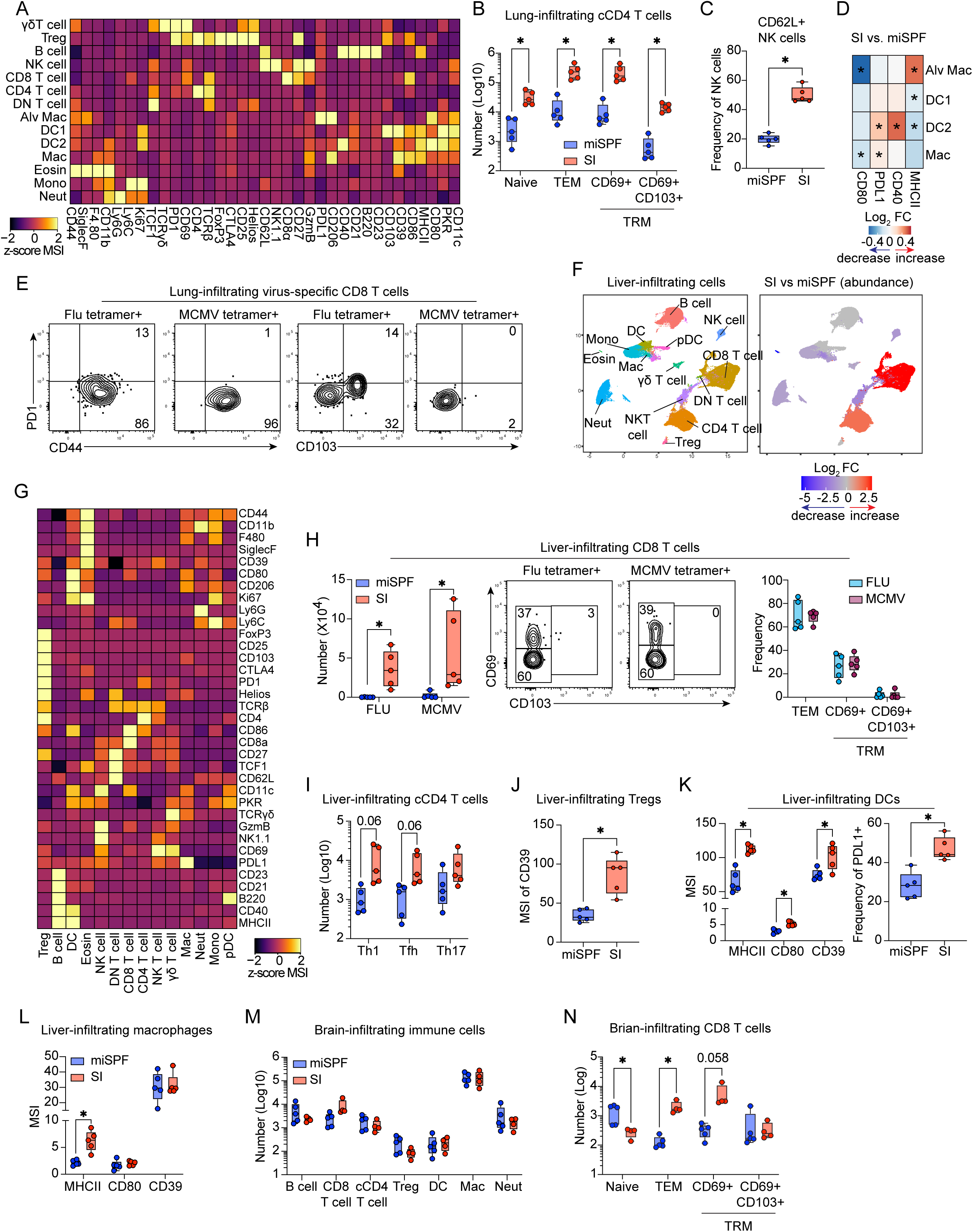
Sequential infection reshapes immune environments in lung, liver, and brain. (A–E) Lung, **(F–L)** liver, and **(M–N)** brain immune landscapes in 18-week-old miSPF and SI mice. **(A)** MSI of phenotypic markers across lung-infiltrating immune subsets. **(B)** Number of lung-infiltrating conventional CD4 T cell subsets. **(C)** Frequency of CD62L expression on lung-infiltrating NK cells. **(D)** Heatmap of log₂ fold change in MSI of MHC II, CD40, PD-L1, and CD80 expression on alveolar macrophages, dendritic cells, and macrophages. **(E)** Expression of PD-1, CD103, and CD44 on lung-infiltrating influenza- or MCMV- tetramer⁺ CD8 T cells. **(F)** UMAP visualization of splenic immune subsets identified by Phenograph clustering. (Left) All immune populations. (Right) Populations with significant changes in abundance, determined using diffcyt. Red: increased in SI mice; blue: decreased in SI mice; gray: unchanged. **(G)** MSI of phenotypic markers across liver-infiltrating immune subsets. **(H)** Representative gating and number of influenza- and MCMV-specific CD8 T cells in liver. **(I)** Number of liver-infiltrating CD4 T helper subsets. **(J)** MSI of CD39 expression on liver Treg cells. **(K)** MSI of MHC II, CD80, and CD39, and frequency of PD-L1 expression on hepatic dendritic cells. **(L)** MSI of MHC II, CD39, and CD80 expression on hepatic macrophages. **(M)** Number of brain-infiltrating immune cell subsets. **(N)** Number of brain-infiltrating CD8 naïve, TEM, and TRM cells. Data represent at least two independent experiments (≥4 mice/group). Circles in the box and whisker plots represent individual mice. Error bars in whisker plots indicate the highest and lowest value, and the line in the box indicates the mean. Statistical significance determined by unpaired t-test. *, p<0.05.

**Figure S3.**
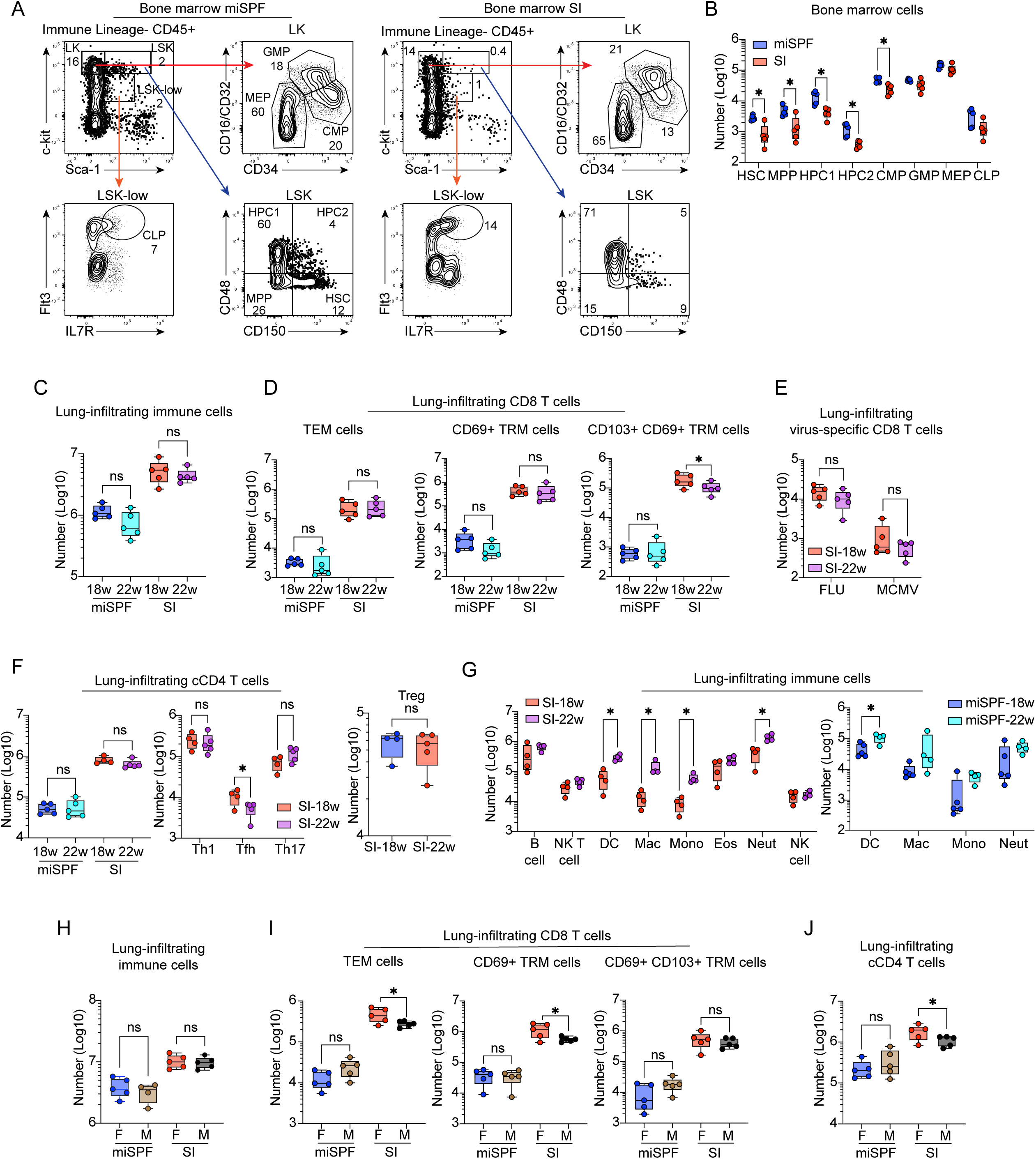
Sequential infection alters hematopoiesis and induces durable, sex-independent immune remodeling. **(A, B)** Bone marrow hematopoietic stem and progenitor cells in 18-week-old female miSPF and SI mice. **(A)** Flow cytometry gating strategy identifying hematopoietic stem and progenitor populations. **(B)** Numerical quantification of hematopoietic stem and progenitor populations. HSC (hematopoietic stem cell), MPP (multipotent progenitor), HPC (hematopoietic progenitor cell), CMP (common myeloid progenitor), GMP (granulocyte-macrophage progenitor), MEP (megakaryocyte-erythroid progenitor), CLP (common lymphoid progenitor). **(C–G)** Lung immune landscape in 18- and 22-week-old female miSPF and SI mice. **(C)** Number of lung-infiltrating (IV-CD45⁻) immune cells. **(D)** Number of lung-infiltrating CD8 TEM and TRM cell subsets. **(E)** Number of lung-infiltrating influenza-specific and MCMV-specific CD8 T cells. **(F)** Number of lung-infiltrating conventional CD4 T cells, T helper subsets, and Tregs. **(G)** Number of lung immune subsets. **(H–J)** Lung immune landscape in 18-week-old male and female miSPF and SI mice. **(H)** Number of lung-infiltrating immune cells. **(I)** Number of lung-infiltrating CD8 TEM and TRM cell subsets. **(J)** Number of lung-infiltrating conventional CD4 T cells. Bone marrow data represent two independent experiments (≥4 mice/group). Experiment comparing male to female, and 18 week to 22 week was performed once with five mice per group. Circles in the box and whisker plots represent individual mice. Error bars in whisker plots indicate the highest and lowest value, and the line in the box indicates the mean. For comparisons in (C)-(G), within SI/miSPF group 18 week was compared to 22 week; for (H)-(J), within SI/miSPF group, female mice were compared to male mice. Statistical significance determined by unpaired t-test. *, p<0.05.

**Figure S4.**
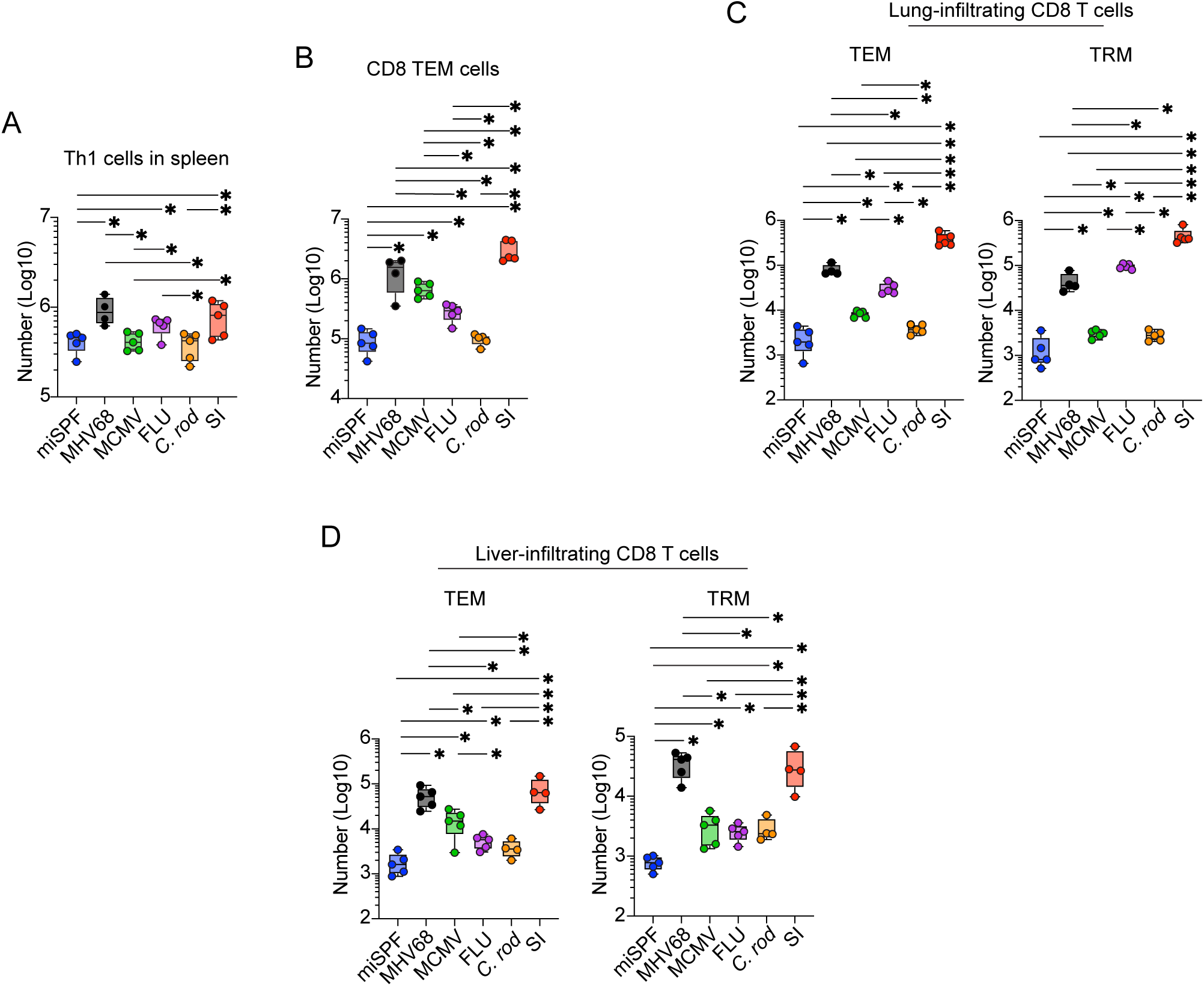
Comparison of SI mice with single infection models. **(A)** Number of splenic CD4 Th1 cells. **(B)** Number of splenic CD8 TEM cells. **(C)** Number of lung-infiltrating CD8 T cell subsets. **(D)** Number of liver-infiltrating CD8 T cell subsets. Data represent two experiments with 4-5 mice per group. Circles in the box and whisker plots represent individual mice. Error bars in whisker plots indicate the highest and lowest value, and the line in the box indicates the mean. Statistical significance determined by unpaired t-test. *, p<0.05.

**Figure S5.**
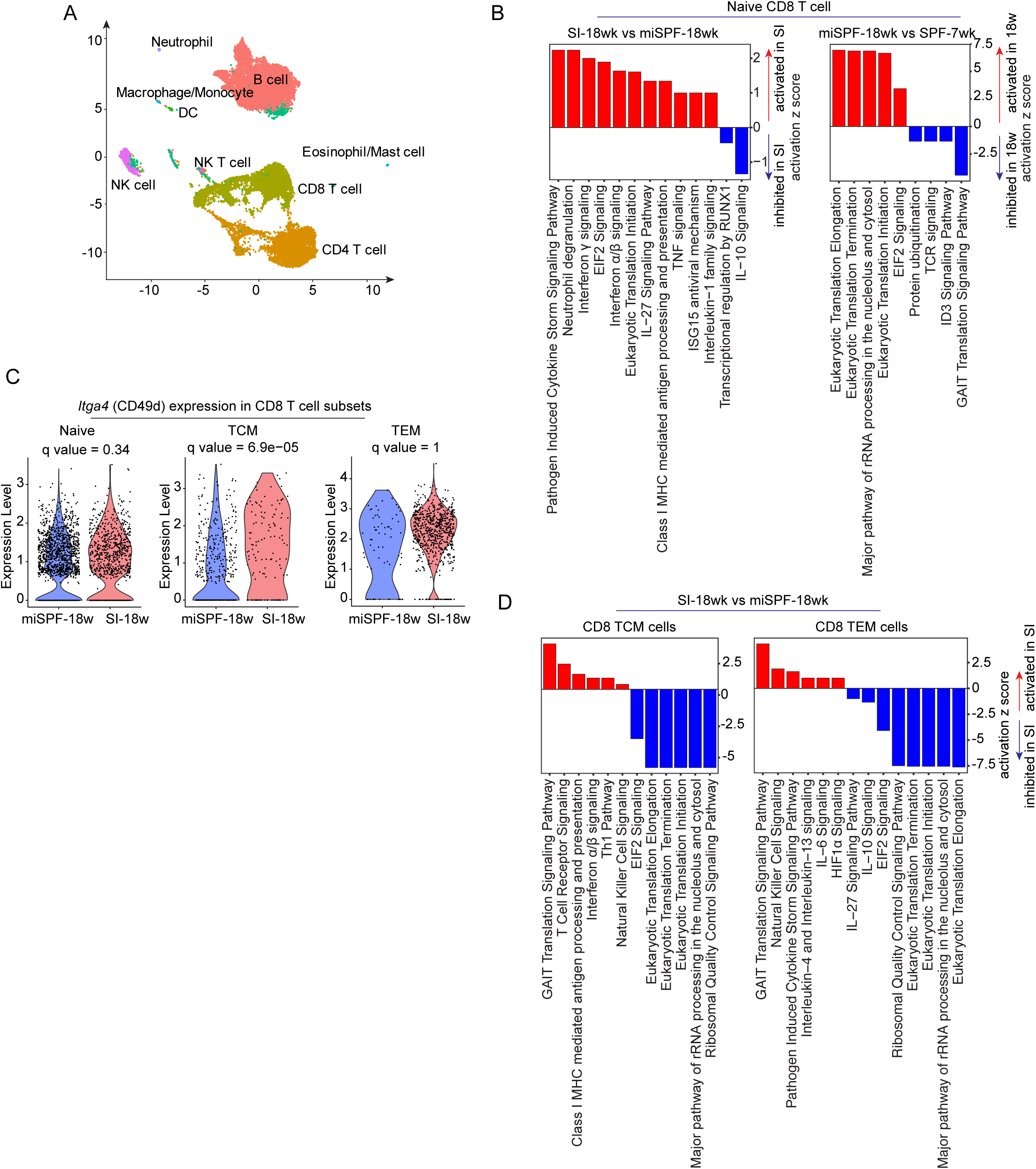
Transcriptomic profiling of splenic immune cells. **(A–D)** scRNA seq of splenocytes from 18-week-old miSPF and SI mice and 7-week-old SPF controls. **(A)** UMAP visualization of all immune cell clusters across conditions. **(B, D)** Pathway analysis of naïve, TCM, and TEM CD8 T cells in 18 week old SI vs 18 week old miSPF and in 18 week old miSPF vs 7 week old SPF mice. **(C)** *Itga4* (CD49d) expression by naïve, TCM, and TEM CD8 T cell subsets.

**Figure S6.**
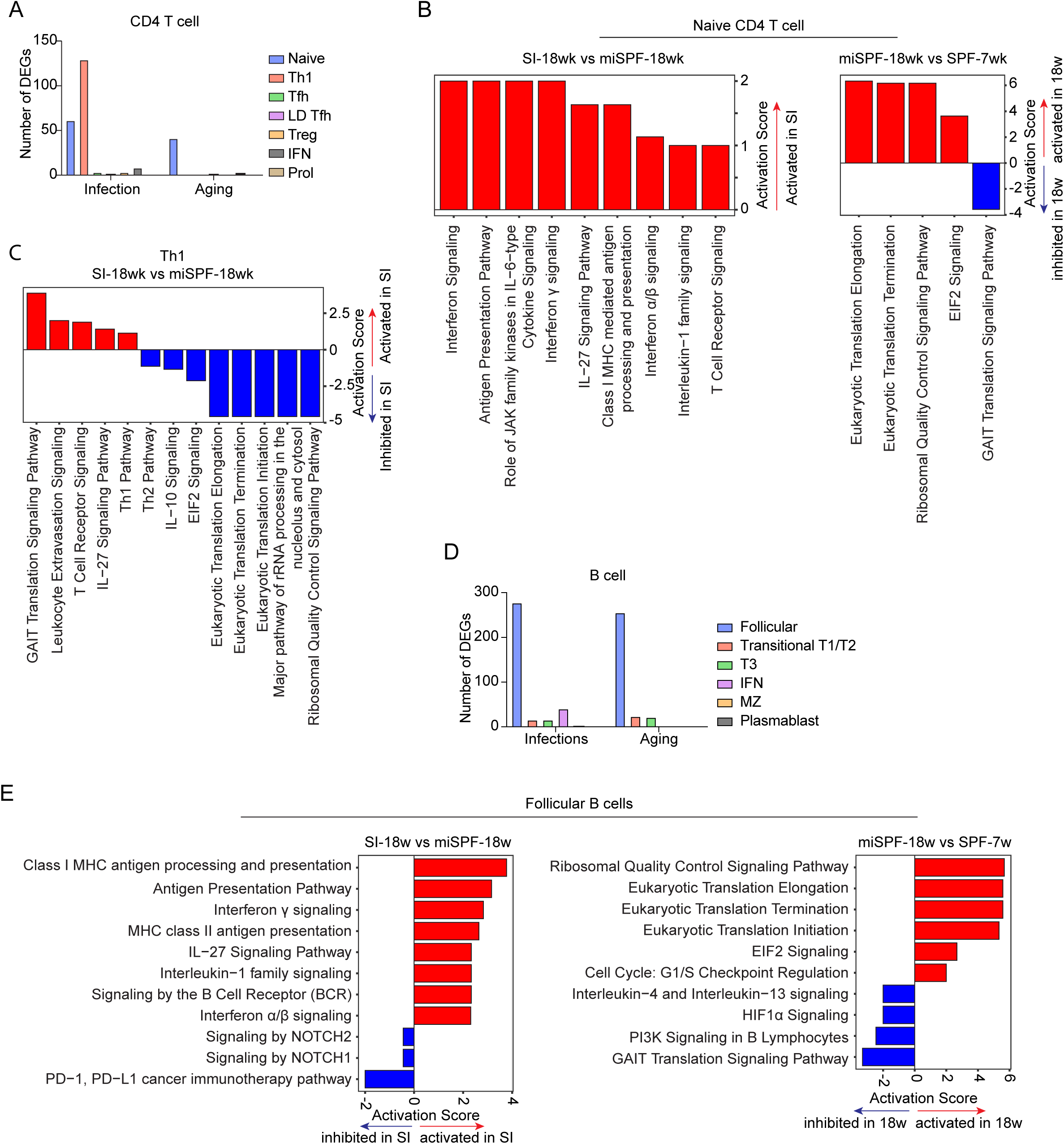
Sequential infection induces transcriptional remodeling of splenic CD4 T cells and B cells. **(A)** Number of DEGs in CD4 T cell subsets attributed to infection or aging. **(B)** Pathway analysis of naïve CD4 T cells comparing 18 week old SI vs 18 week old miSPF and in 18 week old miSPF vs 7 week old SPF mice. **(C)** Pathway analysis of CD4 Th1 cells comparing 18 week old SI vs 18 week old miSPF. **(D)** Number of DEGs in B cell subsets attributed to infection or infection. **(E)** Pathway analysis of follicular B cells comparing 18 week old SI vs 18 week old miSPF and in 18 week old miSPF vs 7 week old SPF mice.

**Figure S7.**
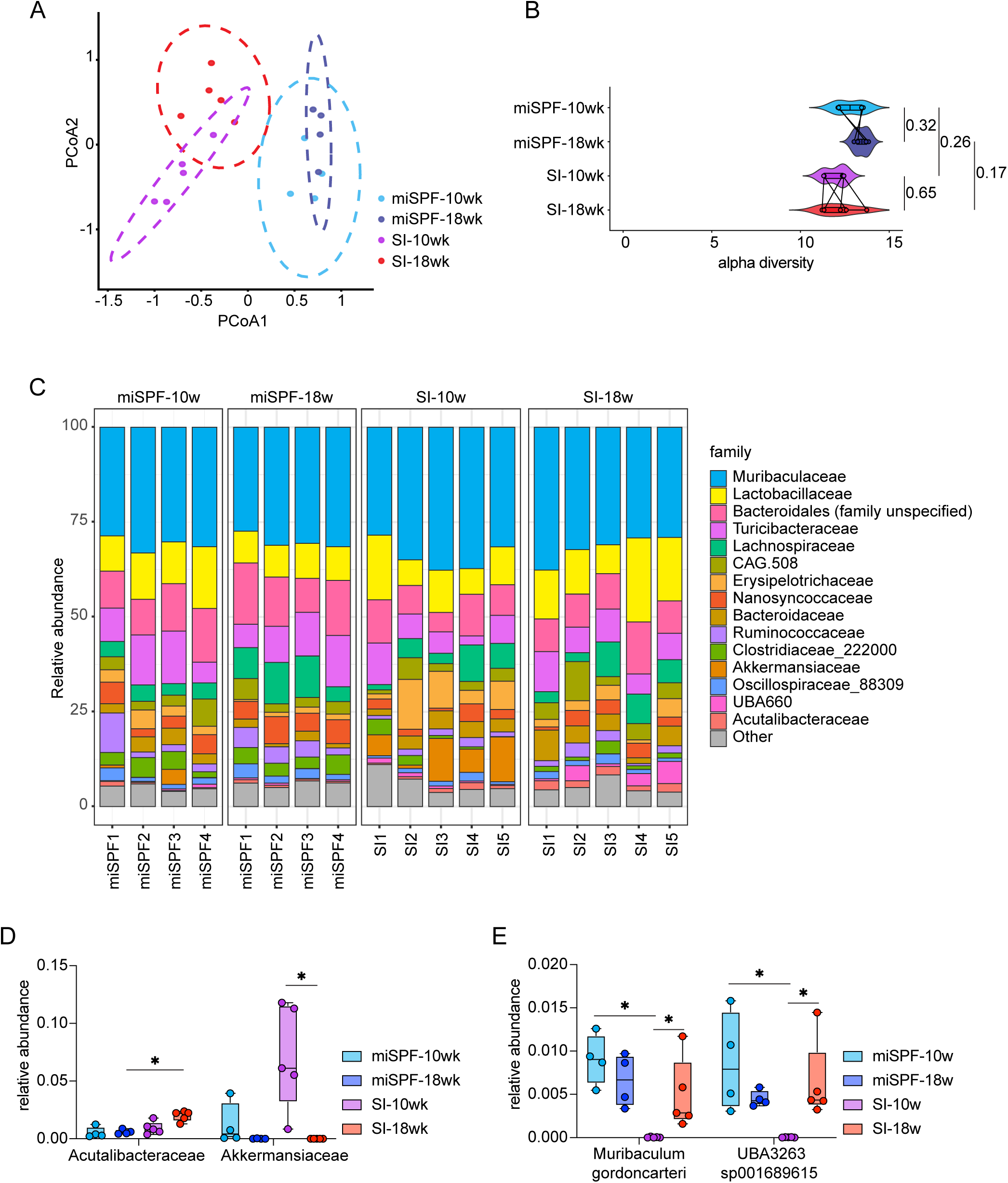
Sequential infection induces modest and transient changes in gut microbiota. (A–E) 16S rRNA sequencing analysis of gut microbiota from miSPF and SI mice at 10 and 18 weeks of age. **(A)** Principal coordinates analysis (PCoA) of microbial community composition; ellipses indicate 95% confidence intervals. **(B)** Faiths PD Alpha diversity measurements. Numbers show the p-adjusted values of pairwise t-test. **(C)** Relative abundance of bacterial families in the stool of individual miSPF or SI mice (columns) at 10 weeks or 18 weeks. The same mice were longitudinally sampled at each time point. **(D)** Relative abundance of bacterial families across comparisons. **(E)** Relative abundance of bacterial species across conditions. Data represent one experiment with 4-5 mice per group. Circles in the box and whisker plots represent individual mice. Error bars in whisker plots indicate the highest and lowest value, and the line in the box indicates the mean. Statistical significance of panel D and E were determined by differential analysis as described in the method section. *, q<0.05

**Figure S8.**
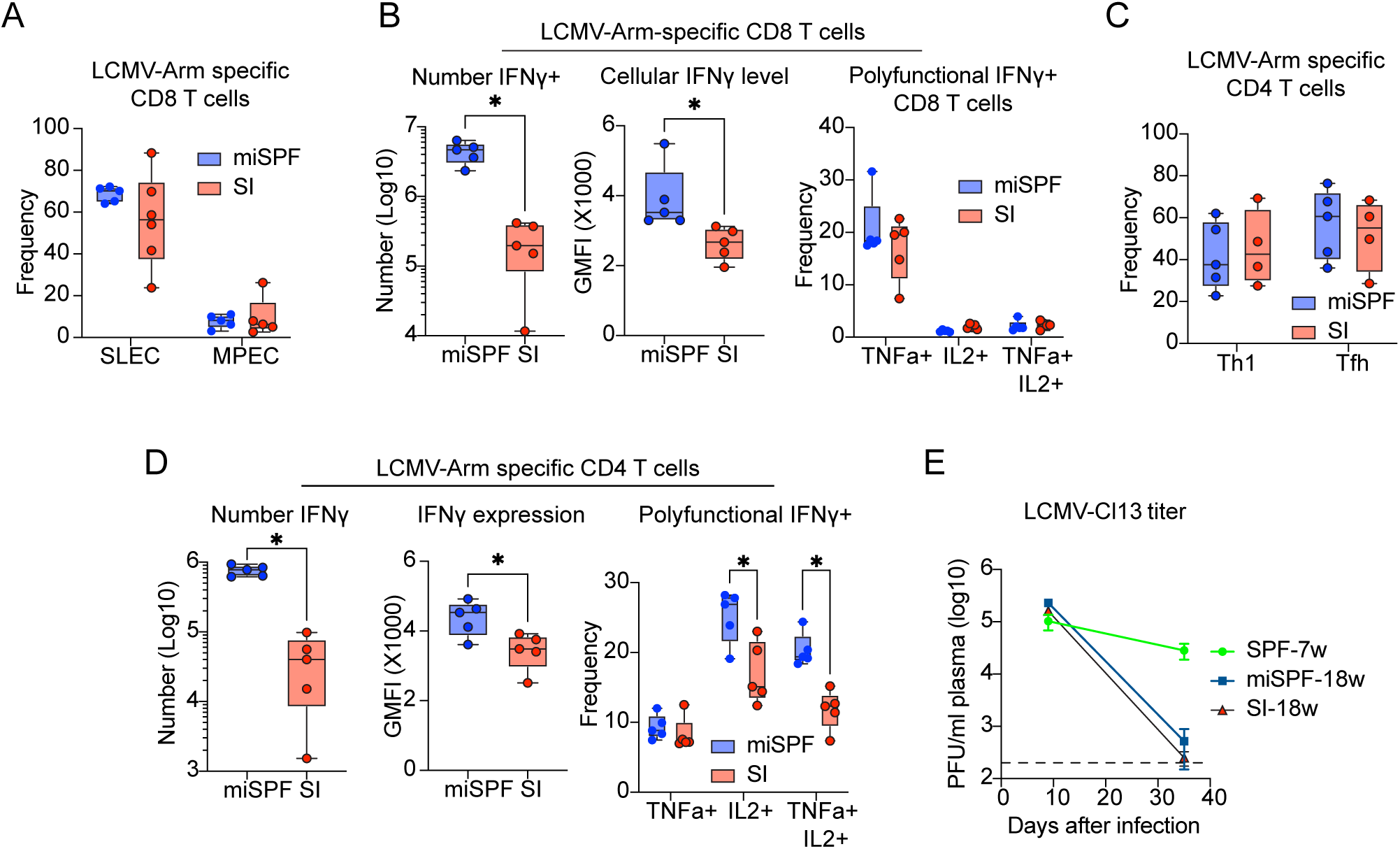
Sequential infection alters immune response to subsequent LCMV infections. (A)-(D) Splenic responses to acute LCMV-Arm infection on day 9 after infection. Characterization of splenic LCMV-GP_33-41_-specific CD8 T cells **(A)** short-lived effector cells (SLECs) and memory precursor effector cells (MPECs) differentiation, **(B)** cytokine production following ex vivo peptide restimulation; and LCMV-GP_61-80_-specific CD4 T cells **(C)** Th1 and Tfh differentiation, **(D)** cytokine production following ex vivo peptide restimulation. **(E)** LCMV-Cl13 titer in the plasma of SPF-7week, miSPF-18week and SI-18week mice. Data represent at least two experiments with 4-5 mice per group, except for cytokine restimulation at day 9 following LCMV-Arm infection, which was only done once with 5 mice per group. Circles in the box and whisker plots represent individual mice. Error bars in whisker plots indicate the highest and lowest value, and the line in the box indicates the mean. Statistical significance determined by unpaired t-test when comparing within groups and one-way ANOVA when also comparing between groups. *, p<0.05.

